# Nanobody-targeted conditional antimicrobial therapeutics

**DOI:** 10.1101/2024.02.20.580917

**Authors:** Chayanon Ngambenjawong, Henry Ko, Novalia Pishesha, Hidde L. Ploegh, Sangeeta N. Bhatia

## Abstract

Conditional therapeutics that rely on disease microenvironment-specific triggers for activation are a promising strategy to improve therapeutic cargos. Among the investigated triggers, protease activity is used most often, because of its dysregulation in several diseases. How to optimally fine-tune protease activation for different therapeutic cargos remains a challenge. Here, we designed nanobody-targeted conditional antimicrobial therapeutics to deliver a model therapeutic peptide and protein to the site of bacterial infection. We explored several parameters that influence proteolytic activation. We report the use of targeting nanobodies to enhance the activation of therapeutics that are otherwise activated inefficiently, despite extensive optimization of the cleavable linker. Specifically, pairing of Ly6G/C or ADAM10-targeting nanobodies with ADAM10-cleavable linkers improved activation via proximity-enabled reactivity. More broadly, this optimization framework provides a guideline for the development of conditional therapeutics to treat various diseases where protease activity is dysregulated.

## Introduction

Most investigational therapeutics fail in clinical trials because they lack efficacy or show unacceptable toxicity.^1–4^ Therapeutics with improved properties are identified through modification of their molecular properties and screening.^5,6^ Formulation of therapeutics with various drug delivery systems (e.g. encapsulation into, or conjugation to nanocarriers or biomacromolecules) may also lead to improvement in efficacy.^7^ Activatable or conditional therapeutics are a promising therapeutic modality to enhance the efficacy or the safety profile of small-molecule drugs and biologics.^8–11^ Specific triggers (e.g. pH, hypoxia, redox, reactive oxygen species, and enzymes) at the diseased sites are exploited to release or convert pro-therapeutics into their active forms. Protease aberration/involvement in the pathology of various diseases has made protease-activated therapeutics a major focus for the development of conditional therapeutics, both pre-clinically and clinically.^8,9,12^

For their activation, several conditional therapeutics rely on passive targeting to engage extracellular proteases at the site(s) of disease.^8,9,12^ Selection of suitable protease-cleavable linkers ensures sufficient on-target activation at the site(s) of disease, with minimal off-target activation. Nonetheless, strategies for the selection and validation of such linkers are still relatively limited. The incorporation of active targeting domains may improve conditional therapeutics, but how this influences accumulation and activation of conditional therapeutics at the site(s) of disease is underexplored. In addition, how different drug carrier compositions affect the activation of a conditional therapeutic is not known.

Here, we report the development and optimization of nanobody (VHH)-targeted conditional antimicrobial therapeutics for conditional delivery of therapeutic peptide and protein cargos to the site of bacterial infection. We systemically investigated the effect of each domain component on conditional activation of the therapeutics. In a mouse model of *Pseudomonas aeruginosa* infection of the lung, we found that Ly6G/C- or ADAM10-targeting VHHs could be used to enhance conditional activation of the tethered therapeutic peptide/protein at the site of infection when paired with ADAM10-cleavable linkers. This enhanced activation was attributed to closer proximity between VHH-targeted therapeutics and the target protease (ADAM10), rather than to increased therapeutic accumulation in the infected organ. We provide a framework for systematic optimization of targeted conditional therapeutics, applicable to diverse disease areas, especially those that involve upregulation of ADAM10, such as infection and cancer.

## Results and discussion

### Design, synthesis, and mechanistic concept of VHH-targeted conditional antimicrobial therapeutics

We previously reported the development of conditional antimicrobial peptide (AMP) therapeutics based on albumin-binding domain (ABD)-AMP conjugates. These conjugates lack a targeting moiety, passively accumulated at the site of bacterial lung infection, and were selectively activated by dysregulated proteases in the affected microenvironment.^13^ Given the clinical efficacy of tumor-targeting antibody-drug conjugates (ADCs) in oncology,^14–16^ antimicrobial therapeutics might similarly benefit from an active targeting domain. Here, we investigated the effect of VHH-mediated targeting on the biodistribution and efficacy of the conditional therapeutics, using both therapeutic peptide and protein cargos. Our targeted conditional antimicrobial therapeutics are composed of (1) a VHH-based active targeting domain, (2) an ABD,^17^ (3) an anionic block/solubility-enhancing domain (EEG)_6_,^13^ (4) a protease-cleavable linker (Sx), and (5) the therapeutic payload (Figure 1A). We used two model cargos (LptD inhibitor POL7080/murepavadin (POL) as a therapeutic peptide and pyocin S2 N-terminal domain-T4 lysozyme (PNT4) as a therapeutic protein).

**Figure 1.**
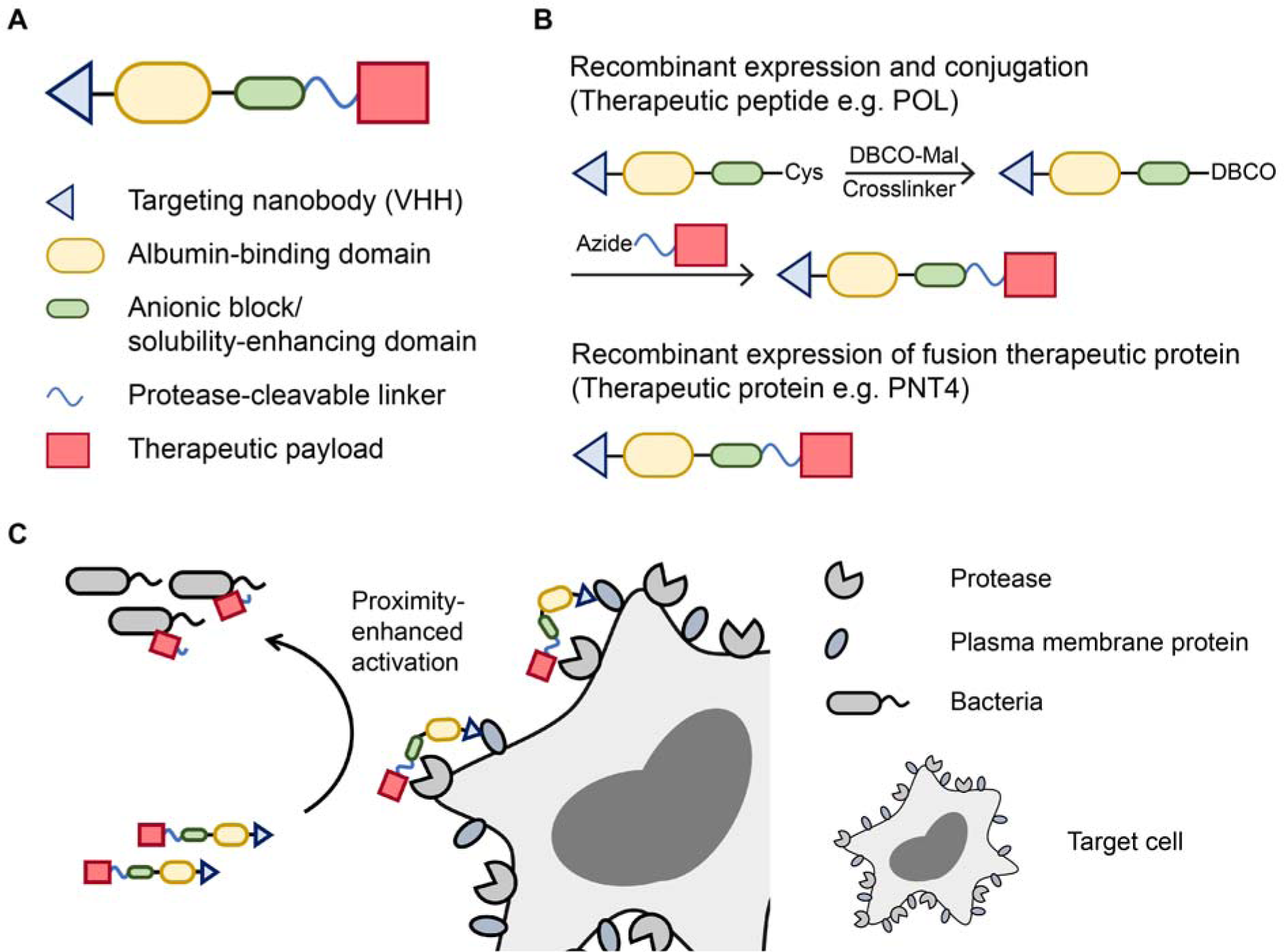
Design, synthesis, and mechanistic concept of VHH-targeted conditional antimicrobial therapeutics. (A) Design and (B) Synthesis of VHH-targeted conditional antimicrobial therapeutics. (C) Proposed mechanism of enhanced conjugate activation via VHH targeting.

The conditional antimicrobial therapeutics were synthesized via (1) conjugation of the VHH-ABD carrier to the peptide cargo or (2) recombinant expression of the full fusion protein combining the VHH-ABD carrier with the therapeutic protein cargo (Figure 1B). For the former, the VHH-(ABD)_2_-(EEG)_6_-Cys carrier domain was recombinantly expressed in *E. coli*, selectively functionalized with dibenzocyclooctyne-maleimide (DBCO-Mal) at the C-terminal Cys, and conjugated to an azide-functionalized cleavable linker-POL fusion peptide (Table S1) via a copper-free azide-alkyne cycloaddition. The fusion peptide was synthesized separately via a standard solid phase-peptide synthesis (SPPS).

We investigated VHHs whose binding targets (e.g. Ly6G/C, CD11b, ADAM10, and ADAM17) are enriched at the site of infection.^18–20^ As model, we used *Pseudomonas aeruginosa* strain O1 (PAO1) to establish a pulmonary infection in mice. We hypothesized that active targeting via VHHs might enhance interaction between infection-localized proteases and the cleavable linker by a proximity effect and so increase activation of the conditional antimicrobial therapeutics (Figure 1C).

### Active targeting with Ly6G/C-targeting VHH16 enhances conditional antimicrobial therapeutic activation

Given the rapid influx of neutrophils at the site of infection,^18^ we first evaluated VHH16, a nanobody that recognizes Ly6G/C, present primarily on monocytes and neutrophils.^21,22^ The schematic of our POL-based targeted antimicrobial conjugate VHH16-(ABD)_2_-(EEG)_6_-S17-POL is shown in Figure 2A. We previously reported an *in vivo* screening pipeline of cleavable linkers in PAO1-infected lungs using ABD-AMP conjugates (ABD)_2_-(EEG)_6_-Sx-(D)Pex-Cy7, where (D)Pex-Cy7 served as a model fluorescently-labeled linear AMP.^13^ In this study, we performed 3 additional rounds of *in vivo* cleavable linker screening (Linker S1-S17) and identified S17 as an improved substrate (Figure S1A-D). This substrate is cleaved by ADAM10/17.^23^ While (ABD)_2_-(EEG)_6_-S17-(D)Pex-Cy7 was efficiently activated in the infected lungs (Figure S1D), the POL conjugate analog (ABD)_2_-(EEG)_6_-S17-POL-Cy7 was poorly activated, possibly due to the increased steric bulk of cyclic peptides (Figure S1E). The need to improve activation of the POL conjugate suggested the inclusion of the VHH targeting domain. We first verified that fluorescently labeled VHH16 accumulated preferentially in PAO1-infected lungs in an infection-dependent, VHH-specific manner (Figure S2). VHH16-(ABD)_2_-(EEG)_6_-S17-POL was readily synthesized. Its identity was confirmed by SDS-PAGE and matrix-assisted laser desorption/ionization-time-of-flight (MALDI-ToF) mass spectrometry (MS) (Figure 2B). The fluorescent version was synthesized using Cy7-labeled POL (Table S1) to enable fluorescent tracking of intact VHH16-(ABD)_2_-(EEG)_6_-S17-POL-Cy7 versus released POL-Cy7, as shown by ADAM10 cleavage (Figure 2C). We confirmed that cleavage of VHH16-(ABD)_2_-(EEG)_6_-S17-POL by ADAM10 conferred conditional activation of the antibacterial activity of POL, showing a 256-fold difference in minimal inhibitory concentrations (MICs) between the intact and activated conjugates (Figure 2D).

**Figure 2.**
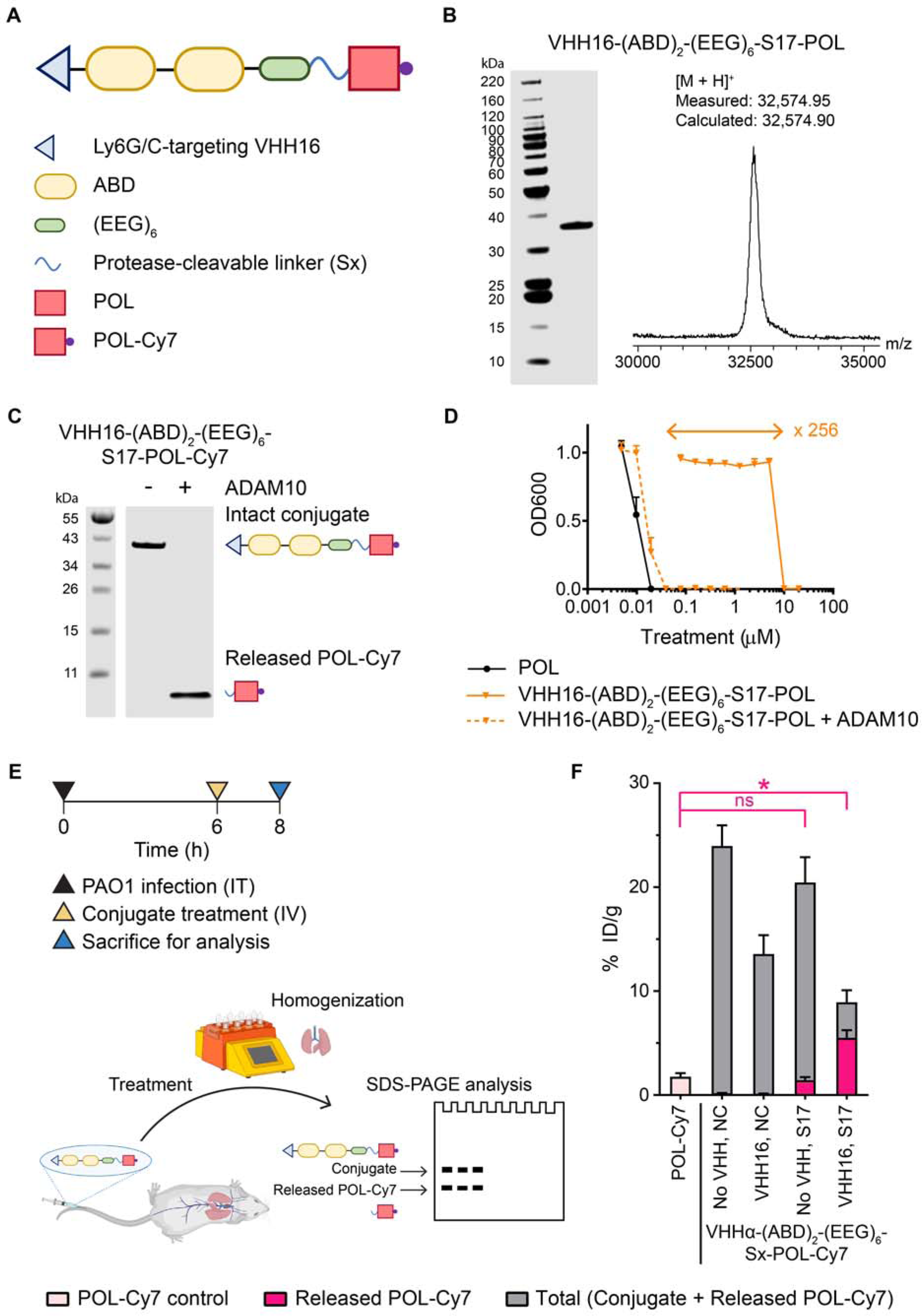
Inclusion of the Ly6G/C-specific VHH16 enhances activation of conditional antimicrobial therapeutic peptide. (A) Design of a conditional antimicrobial therapeutic for delivery of POL. (B) Characterization of POL conjugate VHH16-(ABD)_2_-(EEG)_6_-S17-POL (Left: SDS-PAGE; Coomassie blue staining, Right: Analysis by MALDI-ToF MS reported as mass-to-charge ratio m/z). (C) *In vitro* cleavage assay of VHH16-(ABD)_2_-(EEG)_6_-S17-POL-Cy7 by ADAM10 detected via Cy7 fluorescence using an Odyssey CLx imager. (D) *In vitro* evaluation of masking of antimicrobial activity by VHH16-(ABD)_2_-(EEG)_6_-S17-POL in a microdilution assay on PAO1. Bacterial viabilities were measured based on OD600 absorbance measurements normalized to the untreated control. (E) Experimental timeline and workflow for *in vivo* evaluation of biodistribution and activation of POL-Cy7 conjugates. (F) Quantification of total and activated fractions of the POL-Cy7 conjugates in PAO1-infected lungs presented as % injected dose (ID)/gram (g). Panels D and F were plotted as mean ± standard deviation (SD). (n = 3). Panel F was analyzed with One-way ANOVA with Tukey post hoc tests. Selected comparisons between POL-Cy7 and released POL-Cy7 from the S17 conjugates were shown in pink. * denotes statistical significance (*P* < 0.05). Panel E was partly created with BioRender.com.

To evaluate the biodistribution and activation of VHH16-targeted POL conjugates, non-neutropenic mice were infected intratracheally with PAO1 for 6 h before intravenous treatment with POL-Cy7 conjugates (Figure 2E). Mice were euthanized 2 h after conjugate treatment to harvest and homogenize the infected lungs for subsequent SDS-PAGE quantification of the amounts of the intact conjugate and released POL-Cy7. The non-cleavable and non-VHH control conjugates were included to evaluate both total accumulation and the effect of active targeting. Even though VHH16 itself preferentially accumulated in the PAO1-infected lungs (Figure S2), VHH16-(ABD)_2_-(EEG)_6_-NC-POL-Cy7 with a non-cleavable linker (NC) accumulated approximately 2-fold less compared to non-targeted (ABD)_2_-(EEG)_6_-NC-POL-Cy7 without VHH16 (Figure 2F). A possible explanation could be an on-target, off-infected lung accumulation in other organs. However, VHH16-(ABD)_2_-(EEG)_6_-S17-POL-Cy7 with a cleavable linker S17 was readily activated in the infected lungs, releasing ∼4-fold more POL-Cy7 compared to the amount released from the non-targeted (ABD)_2_-(EEG)_6_-S17-POL-Cy7 (Figure 2F). This enhanced activation of VHH16-(ABD)_2_-(EEG)_6_-S17-POL-Cy7 led to a 3-fold increase in released POL-Cy7 compared to the control (treatment with free POL-Cy7). We confirmed that the albumin-binding half-life extension domain, ABD, is required to achieve the high level of released POL-Cy7 in the infected lungs (Figure S3). Specifically, VHH16-(EEG)_6_-S17-POL-Cy7 without the ABD had equivalent levels of released POL-Cy7 as that of the free POL-Cy7 control in the infected lungs (Figure S3B). The conjugate was cleared via the kidneys (Figure S3D). The roles of active targeting of tumor/infection-targeting ADCs and immunocytokines include increased accumulation in the tumor/infected microenvironment and internalization by cancer/infected cells.^15,24,25^ Here, we report the unique role of such active targeting in the context of protease-activated conditional therapeutics. We find that Ly6G/C-mediated targeting by VHH16 enhances protease activation of its tethered therapeutic conjugate in the infected microenvironment.

### VHH16 enhances activation of conditional antimicrobial therapeutic in a VHH-specific, ADAM10 activity-dependent manner

To better understand the role of active targeting in enhanced activation of conditional antimicrobial therapeutics, we focused on corroborating the identity of the proteases responsible for activation of the therapeutic conjugates. We expanded our screen for cleavable linkers by modification of substrate S17 by truncation or substitutions, as well as by inclusion of an ADAM10-specific linker S26 (TENtide)^23,26^ and a neutrophil elastase (NE)/proteinase 3 (PR3) cleavable linker S27 (Figure 3A). C-terminal truncations of the S17 linker progressively reduced the activation of VHH16-(ABD)_2_-(EEG)_6_-Sx-POL-Cy7 in PAO1-infected lungs. The deletion of C-terminal Leu-Lys (S19) led to a near complete loss of conjugate activation (Figure 3B). This result corresponds to the *in vitro* cleavage assay of the conjugates with recombinant human ADAM10 (Figure S4) and aligns with the reported ADAM10/17 cleavage site between P1(Ala) and P1’(Leu).^23^ P5(Pro-to-Gly), P2(Glu-to-Ala), and P2’(Lys-to-Thr/Val/Ala/Ser) substitutions did not affect the extent of conjugate activation. Even though P2’ and P1’-P2’ truncations are poorly tolerated, flexibility at the P2’ position allows the selection of cleavable linkers whose cleavage scars minimally affect the antimicrobial activity of the tethered therapeutics. For the POL conjugates, the released POL with P1’(Leu)-P2’(Thr) cleavage scar (e.g. cleavage from the linker S22) had an antimicrobial potency equivalent to that of the free POL, whereas the released POL with the P1’(Leu)-P2’(Lys) cleavage scar (from the initial S17 linker) had a 2-fold lower potency (Figure 2D and S5). The conjugate with the TENtide linker S26 was activated as efficiently as the one with the S17 linker, suggesting a contribution of ADAM10 in the infected lungs (Figure 3B). Even though NE is abundant in the infected microenvironment^18^ and the linker S17 can be cleaved by NE *in vitro* (Figure S6), the conjugate with a different NE-cleavable linker (S27),^27^ which is insensitive to ADAM10 cleavage (Figure S4 and S6), was not efficiently activated in the infected lungs. This points to a modest contribution, if any, of NE to *in vivo* activation of the VHH16-based conditional therapeutics (Figure 3B).

**Figure 3.**
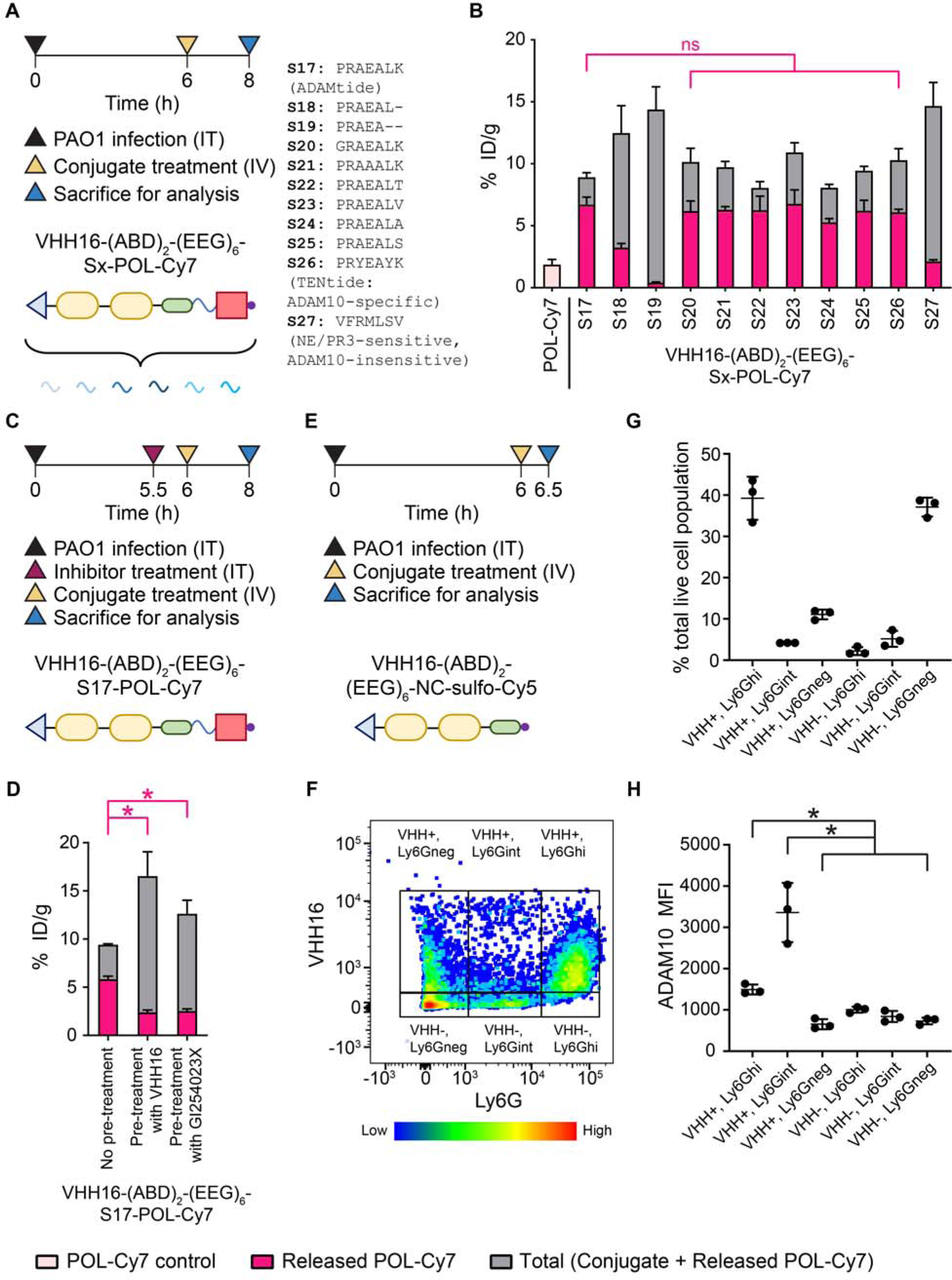
Enhanced activation of a VHH16-targeted conditional antimicrobial therapeutic requires interaction with Ly6G/C target as well as proteolytic activity of ADAM10. (A) Experimental timeline for *in vivo* evaluation of biodistribution and activation of VHH16-(ABD)_2_-(EEG)_6_-Sx-POL-Cy7 with different cleavable linkers (Sx). (B) Quantification of total and activated fractions of the POL-Cy7 conjugates in PAO1-infected lungs presented as % ID/g. (C) Experimental timeline for *in vivo* evaluation of biodistribution and activation of VHH16-(ABD)_2_-(EEG)_6_-S17-POL-Cy7 using intratracheal pre-treatment with either an excess of VHH16-(ABD)_2_-(EEG)_6_ or of the ADAM10-selective inhibitor GI254023X. (D) Quantification of total and activated fractions of VHH16-(ABD)_2_-(EEG)_6_-S17-POL-Cy7 in PAO1-infected lungs presented as % ID/g. (E) Experimental timeline for analysis by flow cytometry of VHH16-(ABD)_2_-(EEG)_6_-NC-SulfoCy5 accumulation in different cell populations of PAO1-infected lungs. (F) A representative dot plot of the lung cell populations based on *in vivo* accumulated VHH16 and *ex vivo* stained Ly6G using an anti-Ly6G monoclonal antibody. Gates were set based on the intensity of VHH16 (+/-) and Ly6G (hi/int/neg). (G) Quantification of each cell population presented as percentage of the total live cell population. (H) Quantification of ADAM10 in each cell population based on *ex vivo* staining with an anti-ADAM10 monoclonal antibody, presented as median fluorescence intensity (MFI). Panels B, D, G, and H were plotted as mean ± SD. (n = 3). Panels B, D, and H were analyzed with One-way ANOVA with Tukey post hoc tests. Selected comparisons between released POL-Cy7 from the S17 conjugate and the other conjugates were shown in pink. * denotes statistical significance (*P* < 0.05).

A follow-up biodistribution study was performed to investigate the effects of binding site saturation or inhibition of ADAM10 on the activation of VHH16-(ABD)_2_-(EEG)_6_-S17-POL-Cy7 *in vivo*. We either administered intratracheally VHH16-(ABD)_2_-(EEG)_6_ (10 eq.) or an ADAM10-selective inhibitor GI254023X, 30 min prior to intravenous treatment with VHH16-(ABD)_2_-(EEG)_6_-S17-POL-Cy7 (Figure 3C). Both lung-localized pre-treatments reduced activation of VHH16-(ABD)_2_-(EEG)_6_-S17-POL-Cy7, confirming a requirement for both VHH16-Ly6G/C-driven interactions as well as ADAM10 catalytic activity in the infected microenvironment (Figure 3D). ADAM10 exists in both transmembrane and soluble forms. We hypothesized that Ly6G/C and ADAM10 might be co-expressed on the same cell population, such that binding of VHH16 brings the conjugate closer to transmembrane ADAM10 and results in proximity-enhanced activation of the conjugate. We therefore performed flow cytometry to evaluate the biodistribution of VHH16-(ABD)_2_-(EEG)_6_-NC-Sulfo-Cy5 in different lung cell populations. VHH16-(ABD)_2_-(EEG)_6_-NC-Sulfo-Cy5 was given intravenously to PAO1-infected mice, followed by euthanasia to obtain lung single cell suspensions for flow cytometry with anti-Ly6G and anti-ADAM10 antibodies (Figure 3E). Gates set based on positivity for VHH16-(ABD)_2_-(EEG)_6_-NC-Sulfo-Cy5 and Ly6G separated lung cell suspension into 6 distinct populations, including VHH16+/VHH16- and Ly6Ghi (high)/int (intermediate)/neg (negative) (Figure 3F). VHH16 accumulated primarily in the Ly6G^hi^ population, and to a lesser extent in the Ly6G^neg^ and Ly6G^int^ populations, respectively (Figure 3F and G). When analyzing these populations for expression of ADAM10, the VHH16^+^Ly6G^int^ population showed the highest level of ADAM10, followed by the VHH16^+^Ly6G^hi^ population. The other cell populations express little, if any, ADAM10 (Figure 3H). Our flow cytometry study confirms co-expression of Ly6G and ADAM10 on a subset of infected lung cell suspension that could therefore be targeted by VHH16, thus supporting our hypothesis of proximity-enhanced activation of such conjugates. Expression of ADAM10 on monocytes/neutrophils has been reported and shown to be necessary for their migration into the alveolar space of lipopolysaccharide (LPS)-induced, inflamed lungs.^28^

### Optimal therapeutic effect of VHH-targeted conditional antimicrobial therapeutic requires optimization of both VHH and cleavable linker

Having observed that active targeting via VHH16 improved activation of the conditional antimicrobial therapeutics, we expanded our investigation to include VHHs that recognize other potentially relevant targets (Host: CD11b, ICAM-1, surfactant protein-A (SP-A), ADAM10, and ADAM17, Pathogen: PcrV) (Figure 4A and Table S2). Another Ly6G/C-targeting VHH clone (VHH21)^21,22^ also improved activation of the POL-based conjugate, albeit to a lower extent than the VHH16-based conjugate (Figure 4B). As seen for the Ly6G/C-targeting VHHs, ADAM10-targeting VHHs also enhanced conjugate activation in a clone-dependent manner. The enhanced activation conferred by the ADAM10 VHHs supports the validity of our hypothesis on proximity-enhanced activity. In an *in vitro* ADAM10 cleavage assay, the ADAM10-targeted conjugates also showed increased activation kinetics, although the trend did not track with *in vivo* activation (Figure S7). Specifically, VHH39G1-(ABD)_2_-(EEG)_6_-S17-POL-Cy7 was the conjugate activated most efficiently *in vivo,* but it had the slowest kinetics of *in vitro* cleavage. This finding highlights the importance of *in vivo* validation to evaluate the extent of conjugate activation. VHHs for the other targets investigated (CD11b, ICAM-1, SP-A, and ADAM17) failed to improve conjugate activation (Figure 4B and S8). The selected VHH clones may simply have been unable to display the conjugates in the appropriate geometry for proper proteolytic activation. Since the goal of conditional therapeutic development is selective activation at the site(s) of disease with minimal exposure in off-target organs, we evaluated the extent of conjugate activation of our lead conjugates in liver and kidneys. For both VHH16-(ABD)_2_-(EEG)_6_-S17-POL-Cy7 and VHH483-(ABD)_2_-(EEG)_6_-S17-POL-Cy7, there is approximately 3-fold more release of POL-Cy7 than seen for the free POL-Cy7 treatment control in PAO1-infected lungs (Figure 4B). These levels were at least 2-fold lower in the liver (Figure 4C) and were comparable in the kidneys (Figure 4D). POL, as an intravenous formulation, has failed a phase III clinical trial due to an increased incidence of acute kidney injury.^4^ There is currently no reported *in vivo* mouse model that recapitulates kidney toxicity induced by POL to guide evaluation of toxicity for new formulations. Even though the current conjugates did not show signs of *in vivo* organ toxicity based on blood chemistry (Figure S9), future directions should focus on improvements in design. This must be done to reduce the extent of off-target activation of conjugates in the kidneys. Investigation into co-formulations with additives such as gelofusine or cilastatin might reduce kidney retention and thus yield further improvements.^29,30^

**Figure 4.**
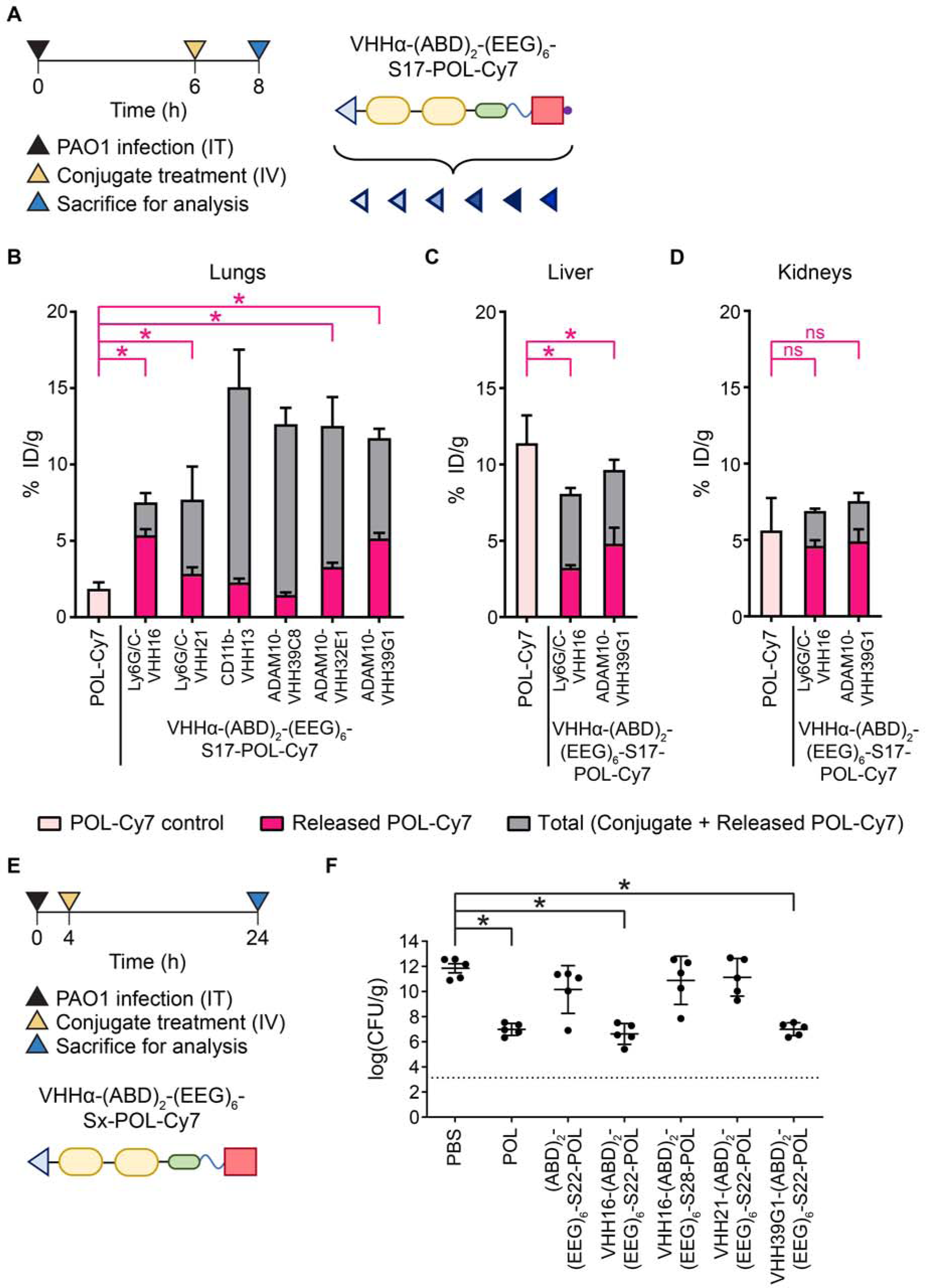
Therapeutic effect of VHH-targeted conditional therapeutic depends on optimization of both VHH and cleavable linker. (A) Experimental timeline for *in vivo* evaluation of biodistribution and activation of VHHα-(ABD)_2_-(EEG)_6_-S17-POL-Cy7 with different targeting VHHs (VHHα). Quantification of total and activated fractions of the POL-Cy7 conjugates in (B) PAO1-infected lungs, (C) liver, and (D) kidneys presented as % ID/g. (E) Experimental timeline for *in vivo* evaluation of therapeutic efficacy of POL conjugates with different targeting VHHs and cleavable linkers. (F) Quantification of bacterial burden from the treated lungs presented as log of colony-forming unit (cfu)/g. Dotted line denotes limit of detection. Panels B-D, and F were plotted as mean ± SD. (n = 3-4 for panel B-D). (n = 5 for panel F). Panels B-D, and F were analyzed with One-way ANOVA with Tukey post hoc tests. Selected comparisons between POL-Cy7 and released POL-Cy7 from the S17 conjugates were shown in pink. * denotes statistical significance (*P* < 0.05). The evaluation of efficacy was confirmed in two independent studies with similar results.

We next evaluated therapeutic efficacy of the conjugates comprising the optimized VHHs together with the optimized cleavable linker. Mice were infected intratracheally with PAO1, followed by treatment with conjugates at 4 h after infection. At 24 h of infection, mice were euthanized to collect the infected lungs for enumeration of bacteria (Figure 4E). Despite the release of more free POL, the lead conjugates (VHH16-(ABD)_2_-(EEG)_6_-S22-POL and VHH39G1-(ABD)_2_-(EEG)_6_-S22-POL) failed to outperform the group that received treatment with free POL (Figure 4F). Both the kinetics of conjugate activation as well as the nature of the acute infection model, which favors immediate high drug bioavailability, may contribute to these findings. Nonetheless, the lead conjugates were more effective than (ABD)_2_-(EEG)_6_-S22-POL, VHH21-(ABD)_2_-(EEG)_6_-S22-POL, and VHH21-(ABD)_2_-(EEG)_6_-S28-POL (NE-specific linker^31^), which match their superior activation *in vivo* (Figure 2F, 4B, and 4F). Our results confirm the need for optimization of conjugate activation to achieve therapeutic efficacy. Future pursuits will examine and optimize yet other parameters that may influence efficacy of the conjugates. The types of infection (e.g. chronic versus acute infection) may matter as well. The drug-release profile of the conjugates described here might result in differential therapeutic efficacy, depending on such contexts. We show the importance of appropriate pairing between active targeting VHHs (Ly6G/C and ADAM10 binders) and cleavable linkers (ADAM10 substrates) to achieve enhanced conjugate activation. The lead VHHs in this study (Ly6G/C and ADAM10 binders) accumulated preferentially in the lungs of mice infected with PAO1 (Figure S2 and S10). Even so, further screens for VHHs that target a more infection-selective cell population (e.g. CD177 on activated neutrophils) or that interact with other dysregulated proteases in the diseased microenvironment (e.g. NE and PR3) may allow further improvement.^32–35^ Exploiting dysregulated proteases in the diseased microenvironment presents an additional avenue for conditional therapeutic development, provided the optimal combination of VHH target and cleavable linker can be identified.

### Active targeting with Ly6G/C-targeting VHH16 enhances conditional activation of a model therapeutic protein

Following optimization of the conditional antimicrobial therapeutic with a therapeutic peptide, POL, we investigated whether this design could be similarly adapted for conditional delivery of a therapeutic protein. With their unique mechanism of action, engineered lysins are a potential alternative to antibiotics.^36,37^ Lysins hydrolyze peptidoglycan, readily killing gram-positive bacteria, but are unable to penetrate the gram-negative bacterial outer membrane and reach the periplasm where peptidoglycan is accessible for bactericidal action. Lysins fused to bacterial internalization domains enable their import into periplasm.^37–39^ We reasoned that by sterically blocking the internalization domain with the VHH-ABD fusion, it would be possible to formulate a conditional antimicrobial therapeutic. Here, we used the N-terminal domain of pyocin S2 (Amino acid 1 - 209) to enable import of T4 lysozyme (Catalytically active, disulfide-free mutant (C54T, C97A)^40^) into PAO1 which expresses the pyocin S2 receptor FvpAI^41^ (Figure 5A). The conditional therapeutic VHH16-ABD-(EEG)_6_-S17-PNT4 was readily expressed as a fusion protein in *E. coli* (Figure S11). The C-terminal sulfo-Cy7-labeled version was used to confirm conditional activation via cleavage by ADAM10 (Figure 5B), which corresponds to masking of conditional bactericidal activity (Figure 5C).

**Figure 5.**
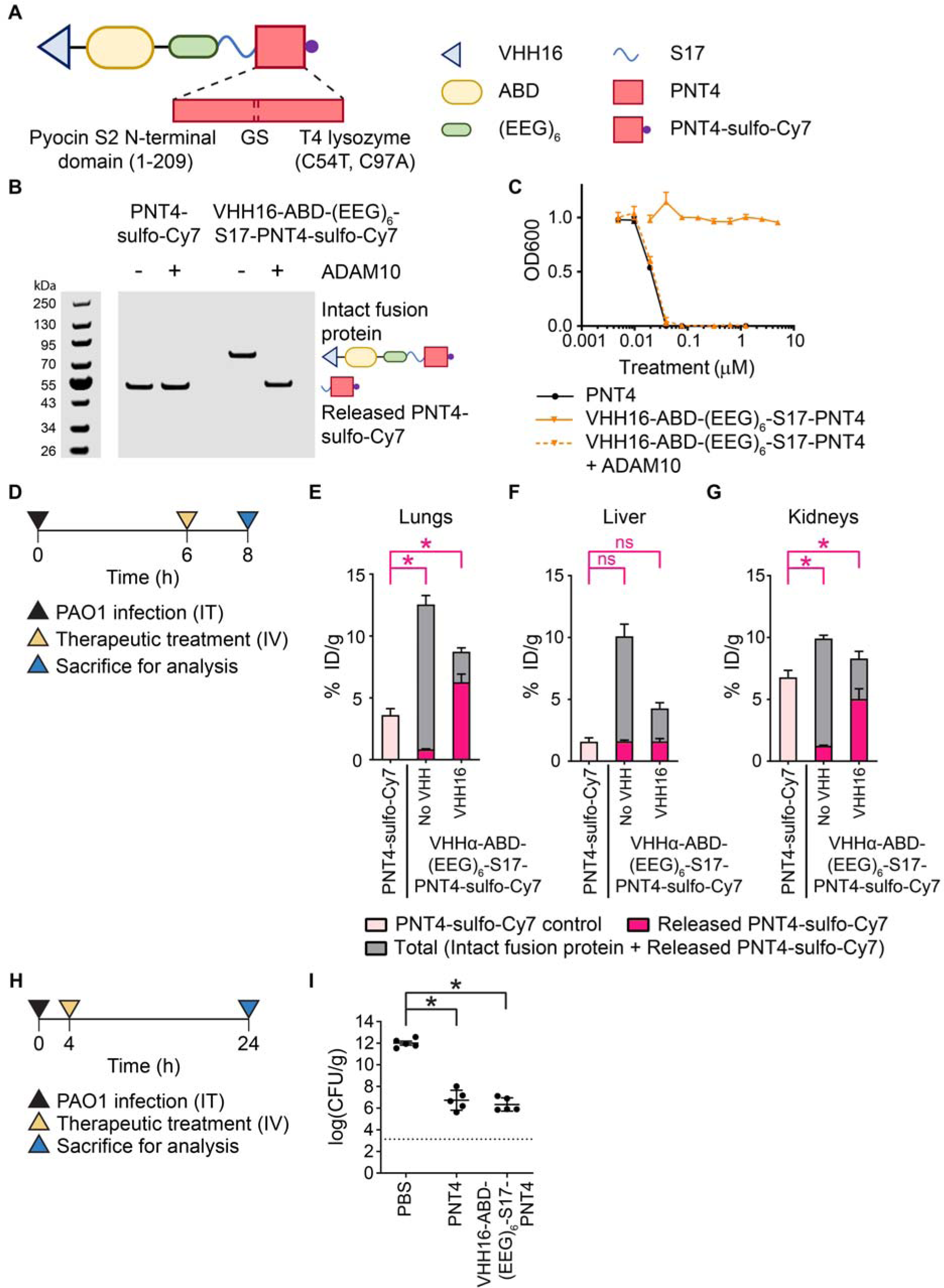
Demonstration of VHH16-enhanced activation of a conditional antimicrobial therapeutic protein. (A) Design of conditional antimicrobial therapeutic for delivery of PNT4. (B) *In vitro* cleavage assay of VHH16-ABD-(EEG)_6_-S17-PNT4-sulfo-Cy7 by ADAM10 detected via sulfo-Cy7 fluorescence using an Odyssey CLx imager. (C) *In vitro* evaluation of antimicrobial activity masking of VHH16-ABD-(EEG)_6_-S17-PNT4 via microdilution assay on PAO1. Bacteria viabilities were measured based on OD600 absorbance normalized to the untreated control. (D) Experimental timeline for *in vivo* evaluation of biodistribution and activation of VHH16-ABD-(EEG)_6_-S17-PNT4-sulfo-Cy7. Quantification of total and activated fractions of the PNT4-sulfo-Cy7 in (E) PAO1-infected lungs, (F) liver, and (G) kidneys presented as % ID/g. (H) Experimental timeline for *in vivo* evaluation of therapeutic efficacy of VHH16-ABD-(EEG)_6_-S17-PNT4. (I) Quantification of bacterial burden from the treated lungs presented as log(cfu/g). Dotted line denotes limit of detection. Panels C, E-G, and I were plotted as mean ± SD. (n = 3 for panels C and E-G). (n = 5 for panel I). Panels E-G, and I were analyzed with One-way ANOVA with Tukey post hoc tests. Selected comparisons between PNT4-sulfo-Cy7 and released PNT4-sulfo-Cy7 from the conditional therapeutics were shown in pink. * denotes statistical significance (*P* < 0.05). The evaluation of efficacy was confirmed in two independent studies with similar results.

A biodistribution study in PAO1-infected mice was performed to compare VHH16-ABD-(EEG)_6_-S17-PNT4-sulfo-Cy7, ABD-(EEG)_6_-S17-PNT4-sulfo-Cy7, and PNT4-sulfo-Cy7 (Figure 5D). As observed for the POL conjugates, the non-targeted ABD-(EEG)_6_-S17-PNT4-sulfo-Cy7 was poorly activated in the infected lungs, whereas the addition of VHH16 (VHH16-ABD-(EEG)_6_-S17-PNT4-sulfo-Cy7) led to a 7.5-fold increase in activation (Figure 5E). VHH16-ABD-(EEG)_6_-S17-PNT4-sulfo-Cy7 released 1.7-fold more active protein in the infected lungs than did treatment with the free PNT4-sulfo-Cy7. In the off-target organs, the active fractions were comparable in the liver (Figure 5F) and 1.4-fold lower for VHH16-ABD-(EEG)_6_-S17-PNT4-sulfo-Cy7 in the kidneys compared to the free PNT4-sulfo-Cy7 group (Figure 5G). Evaluation of *in vivo* efficacy showed comparable potency for VHH16-ABD-(EEG)_6_-S17-PNT4 and free PNT4 (Figure 5H and 5I). It is possible that the fold difference in the active protein fractions is not sufficient to boost therapeutic efficacy in this infection model. Further optimization should be considered similar to what was described for the POL conjugates. PNT4 was well-tolerated in mice due to its unique specificity for the hydrolysis of peptidoglycan (Figure S12). The design of conditional antimicrobial therapeutics and the parameters for optimization could be applied to other therapeutic peptides or proteins where enhanced delivery is beneficial or where off-target toxicity might be a concern.

## Conclusion

We report the development of nanobody-targeted conditional antimicrobial therapeutics for enhanced delivery of therapeutic antimicrobial peptide (POL) and protein (PNT4) to the site of infection. The pairing of VHHs that target Ly6G/C or ADAM10 with ADAM10-cleavable linkers was responsible for improved activation. Analysis by flow cytometry of infected lung cell populations that bound Ly6G/C-specific VHH16 showed a subpopulation that co-expressed both Ly6G and ADAM10. This co-localization supports the postulated mechanism of improved activation via a proximity-enhanced reactivity. The selection of Ly6G/C or ADAM10-specific VHHs as targeting moieties and the optimization of ADAM10 cleavable linkers were essential to improve therapeutic efficacy of these conditional antimicrobial therapeutics. Other targeting domains (e.g. CD177/Ly6G/NE/PR3/MMP8 binders), cleavable linkers (e.g. NE/PR3/MMP8 substrates), and combinations merit consideration to further improve specificity for the infected microenvironment or to improve delivery of the active therapeutic payloads. Our platform and optimization framework are applicable to the development of conditional therapeutics for other disease areas with dysregulated proteases, including cancer, autoimmune disease, and fibrosis.

## Materials and methods

### Molecular cloning

gBlocks^TM^ gene fragments encoding fusion proteins flanked with NcoI and XhoI restriction sites were ordered from Integrated DNA Technologies (IA, U.S.A.). The gene fragments were cloned into a Novogen pET28a(+) plasmid vector at the NcoI and XhoI restriction sites via restriction enzyme digestion and ligation and transformed into NEB® 5-alpha competent *E. coli*. Colonies containing correct sequences of the gene inserts were identified by Sanger sequencing via Quintara Biosciences (MA, U.S.A.) and grown overnight for miniprep extraction of plasmid DNA. The extracted plasmids were transformed into BL21(DE3) competent *E. coli* for non-disulfide bond-containing proteins or SHuffle® T7 Express competent *E. coli*. for disulfide bond-containing proteins (e.g. nanobody-containing fusion proteins).

### Recombinant expression of protein therapeutics

Overnight primary cultures of BL21(DE3) or SHuffle® T7 Express *E. coli* encoding the proteins of interest were expanded into 500-mL secondary culture in Luria-Bertani (LB) broth (supplemented with 50 μg/mL Kanamycin) and incubated in an incubator shaker at 220 rpm and 37 °C for 4-6 h until OD600 reached 0.6-0.8. The BL21(DE3) *E. coli* secondary culture was induced with 1 mM isopropyl β-D-1-thiogalactopyranoside (IPTG) for 2 h at 37 °C while the SHuffle® T7 Express *E. coli* secondary culture was induced with 0.4 mM IPTG for 24 h at 25 °C. At the end of the IPTG induction, bacteria were spun down at 4,500 rpm for 15 min, frozen, and stored in a -80 °C freezer. For purification, bacteria pellet was thawed on a 37 ◦C water bath, resuspended in a B-PER^TM^ complete bacterial protein extraction reagent (15 mL), and incubated on a shaker for 15 min at room temperature (RT). The lyzed bacteria suspension was spun down at 11,000 rpm for 20 min and the supernatant was incubated in a Qiagen Ni-NTA agarose resin (1 mL) for 1 h at 4 °C to capture the His-tagged protein product. The resin suspension was poured into a fritted column, washed with 5 mL of a wash buffer A (50 mM Tris, 500 mM NaCl, 2% Triton X-114), 5 mL of a wash buffer B (50 mM Tris, 500 mM NaCl), and eluted with 1.5 mL of an elution buffer (50 mM Tris, 500 mM NaCl, 500 mM imidazole, 10% glycerol). The eluted product was confirmed by SDS-PAGE using NuPAGE™ 4 to 12% Bis-Tris mini protein gel stained with Bio-Safe^TM^ Coomassie stain.

### Site-specific conjugation of therapeutic peptide or fluorescent dye

Selective reduction of the C-terminal cysteine of VHH-containing fusion protein was adapted from a previously described protocol.^42^ In brief, the protein solution (5-10 mg in 1.5 mL elution buffer) was supplemented with EDTA (1 mM final concentration) and incubated in Pierce^TM^ immobilized TCEP disulfide reducing gel (0.25 mL, pre-washed thrice with 1 mL PBS) with gentle rotation at RT for 24 h. Following the incubation, the gel was pelleted at 5,000 rpm for 5 min. The supernatant was centrifuge-filtered to exchange buffer into PBS (1 mM EDTA, pH 6.5) using 10-kDa Amicon centrifugal filter units (2 filter units per 5 mg protein, 14,000 rpm spin for 2 min, 4 times). The protein solution was diluted in the same buffer to ∼5 mg/mL and reacted with DBCO-Mal (3 eq.) or fluorescent dye-maleimide (1.5 eq.) at RT for 4 h. The DBCO-functionalized or dye-labeled protein was purified using a Cytiva disposable PD-10 desalting column to remove unreacted DBCO-Mal/dye-maleimide and exchange the buffer for PBS (pH 7.4). Therapeutic peptides and their Cy7-labeled analogs (Table S1) were synthesized by CPC Scientific (CA, U.S.A.) via standard Fmoc-based SPPS. For therapeutic peptide conjugation, DBCO-functionalized protein (10 mg) was first immobilized onto Qiagen Ni-NTA agarose (0.5 mL, pre-washed thrice with 1 mL PBS) at RT for 15 min. Azido-functionalized therapeutic peptide (2 eq.) was then added to the agarose suspension and incubated with gentle rotation at RT for 24 h. The agarose suspension was loaded into a fritted column, washed with the wash buffer B (10 mL) to remove unreacted peptide, and eluted with the elution buffer (1 mL). The conjugated product was buffer-exchanged into PBS (pH 7.4) using a PD-10 desalting column and its identity and purity were confirmed by SDS-PAGE. The lead candidates were further characterized by MALDI-ToF MS.

### Protease cleavage assay

Dye-labeled antimicrobial therapeutics (VHHα-(ABD)_2_-(EEG)_6_-Sx-POL-Cy7 or VHH16-ABD-(EEG)_6_-S17-PNT4-sulfo-Cy7) were incubated with recombinant human ADAM10 (R&D Systems, MN, U.S.A.) at 10 μM and 250 nM final concentrations respectively. Following the incubation at RT for 24 h, an aliquot (5 μL) was collected and diluted into PBS (1x Halt protease inhibitor cocktail, 1x EDTA) (15 μL) to stop the protease activity. SDS-PAGE was performed using NuPAGE™ 4 to 12% Bis-Tris mini protein gel to separate cleaved and intact therapeutics and detect via Cy7/sulfo-Cy7 fluorescence using an Odyssey CLx imager at the 800 nm channel (LI-COR Biosciences, NE, U.S.A.).

### Microdilution assay

*P. aeruginosa* strain PAO1 was a generous gift from the Ribbeck Lab at the Massachusetts Institute of Technology. A secondary culture of PAO1 was grown in Mueller Hinton broth (MHB) in an incubator shaker at 37 °C until OD600 reached approximately 0.6. The bacteria culture was washed once in MHB, resuspended in MHB (1 mM human serum albumin (HSA)) to 10^6^ cfu/mL density, and plated on 96-well plates (50 μL/well). Two-fold serial dilutions of antimicrobial therapeutic protein solutions (with or without pre-incubation in human ADAM10 (250 nM) for 8 h) were prepared in MHB and transferred to the bacteria-plated wells to the final volume of 100 μL (5 x 10^5^ cfu per well). The plates were incubated at 37 °C for 16 h before measurement of OD600 using an Infinite 200 PRO plate reader (Tecan, Switzerland) to determine bacteria viability.

### Mouse model of bacterial lung infection

All animal studies were approved by the Massachusetts Institute of Technology’s Committee on Animal Care (MIT CAC protocol 2203000310). For a neutropenic lung infection model, CD-1 mice (11-12 weeks old) were intraperitoneally administered cyclophosphamide at 4 and 1 days (150 and 100 mg/kg respectively) prior to infection. On the infection day, a secondary culture of PAO1 (OD600 ∼0.6-0.8) was washed twice in PBS, resuspended in PBS, and intratracheally administered to mice (2 × 10^5^ cfu in 50 μL PBS) using a 22G blunt-end catheter (EXCEL International). For a non-neutropenic lung infection model, the cyclophosphamide treatment was omitted, and the PAO1 suspension was intratracheally administered at the dose of 7.5 x 10^6^ cfu in 50 μL PBS. All animal studies were performed in the non-neutropenic lung infection model unless specified as the neutropenic model.

### Biodistribution study

After 6 h of bacterial lung infection, the infected mice were intravenously administered dye-labeled antimicrobial therapeutics (15 nmol) and euthanized 2 h later to collect the infected lungs, liver, and kidneys. The organs were transferred to gentleMACS^TM^ M tubes and homogenized on a gentleMACS^TM^ tissue dissociator using PBS (1x Halt protease inhibitor cocktail) as a medium. The homogenates were pelleted at 14,000 rpm for 30 min, and the supernatants were used for analysis. SDS-PAGE was performed using NuPAGE™ 4 to 12% Bis-Tris mini protein gel to separate cleaved and intact therapeutics and detect via Cy7/sulfo-Cy7 fluorescence using an Odyssey CLx imager at the 800 nm channel. Quantification (% ID/g) was determined using a standard curve from the stock solution of the dye-labeled therapeutics. For the biodistribution study with VHH16 competition or ADAM10 protease inhibitor, either VHH16-(ABD)_2_-(EEG)_6_ (10 eq. in 50 μL PBS) or ADAM10-selective inhibitor GI254023X (MedChemExpress, NJ, U.S.A.) (5 mg/kg dose in 50 μL 0.9% NaCl (10% DMSO, 20% sulfobutylether-β-cyclodextrin)) was administered intratracheally at 5.5 h post-infection followed by intravenous treatment with VHH16-(ABD)_2_-(EEG)_6_-S17-POL-Cy7 (10 nmol) at 6 h post-infection.

### Flow cytometry analysis of VHH biodistribution in lungs

After 6 h of bacterial lung infection, the infected mice were intravenously administered VHH16-(ABD)_2_-(EEG)_6_-NC-sulfo-Cy5 without a cleavable linker (10 nmol) and euthanized 30 min later to collect the infected lungs. Single cell suspensions of the infected lungs were prepared as previously described.^43^ Briefly, the infected lungs were transferred to gentleMACS^TM^ C tubes and homogenized on a gentleMACS^TM^ tissue dissociator (“m_lung_01” program) using HEPES buffer (2 mg/ml collagenase D (Roche) and 80 U/mL DNAse I (Roche)) as a medium. The homogenates were incubated in an incubator shaker at 37 °C for 30 min and further homogenized using the “m_lung_02” program. Single cell suspensions were collected by filtering the homogenates through 70 μm mesh cell strainers. The single cell suspensions were centrifuged at 300 g for 10 min and resuspended in PBS (3 mL). Aliquots of the single cell suspensions (5 x 10^6^ cells per aliquot) were first stained with Zombie Aqua™ fixable viability kit (1:500 dilution in PBS) and Fc-blocked with TruStain FcX^TM^ (anti-mouse CD16/32) antibody (BioLegend, CA, U.S.A.) (1:20 dilution in PBS (1% BSA)) before further staining with Alexa Fluor^TM^ 488 anti-mouse Ly6G antibody (Clone 1A8) (BioLegend, CA, U.S.A.) (1:100 dilution in PBS (1% BSA)) and DyLight^TM^ 550 anti-mouse ADAM10 antibody (Clone RM0146-7H12) (Novus Biologicals, CO, U.S.A.) (1:100 dilution in PBS (1% BSA)). The stained cell suspensions were fixed in 4% paraformaldehyde and resuspended in PBS (1% BSA) before flow cytometry analysis using a FACSymphony™ A3 cell analyzer (BD Biosciences, NJ, U.S.A.).

### *In vivo* efficacy evaluation

After 4 h of bacterial lung infection, the infected mice were intravenously administered antimicrobial therapeutics (2.5 mg/kg POL eq. for POL and POL conjugates and 0.25 mg/kg PNT4 eq. for PNT4 and VHH16-ABD-(EEG)_6_-PNT4). At 24 h post-infection, the mice were euthanized. The infected lungs were collected and homogenized on a gentleMACS^TM^ tissue dissociator using gentleMACS^TM^ M tubes with PBS as a medium. The homogenates were ten-fold serially diluted in PBS and plated on LB agar plates to determine cfu/g.

### *In vivo* toxicity evaluation

Healthy CD-1 mice were intravenously administered antimicrobial therapeutics (5 mg/kg POL eq. per dose for POL and POL conjugates (2 doses at 0 and 6 h time points) and 5 mg/kg PNT4 eq. for PNT4 and VHH16-ABD-(EEG)_6_-PNT4) and monitored for weight change and any signs of distress. At 24 h post-treatment, the mice were euthanized via cardiac puncture to draw blood for serum separation using Microtainer^TM^ serum separator tubes. The serum samples were submitted to the MIT Division of Comparative Medicine Diagnostic Laboratory for serum chemistry analysis.

### Statistical analysis and schematic representation

Statistical analysis was performed on GraphPad Prism software. Data were plotted as mean ± SD. Comparisons among different treatment groups were based on One-way ANOVA with Tukey post hoc tests. *P* value < 0.05 was considered statistical significance. A part of schematics in this publication was created with BioRender.com.

## Associated content

### Supporting information

Figure S1. Optimization of cleavable linker improves conditional therapeutic activation, Figure S2. VHH16 accumulated in PAO1-infected lungs in an infection-dependent, VHH-specific manner, Figure S3. ABD is required to increase the amount of released POL-Cy7 of VHH16-targeted conjugate in PAO1-infected lungs, Figure S4. *In vitro* ADAM10 cleavage assay identified tolerable mutations of the S17 linker, Figure S5. Activated VHH16-(ABD)_2_-(EEG)_6_-S22-POL has an equivalent antimicrobial potency as free POL, Figure S6. S17 and S27 POL-Cy7 conjugates can be activated to neutrophil elastase, Figure S7. ADAM10-targeting VHHs enhance *in vitro* conjugate activation by ADAM10, Figure S8. Expanded screening of VHHs with relevant targets for enhanced conjugate activation, Figure S9. VHH-targeted POL conjugates exhibit good safety profiles, Figure S10. VHH39G1 accumulated in PAO1-infected lungs in an infection-dependent, VHH-specific manner, Figure S11. Recombinant VHH16-ABD-(EEG)_6_-S17-PNT4 was readily expressed, Figure S12. VHH-targeted conditional PNT4 exhibits a good safety profile, Table S1. List of therapeutic peptides and sequences, Table S2. List of VHH clones.

## Acknowledgment

We would like to acknowledge H. Fleming for critical editing of the manuscript. We would like to acknowledge Koch Institute Swanson Biotechnology Center Flow Cytometry Core, especially M. Griffin and G. Paradis for consultation on flow cytometry. We would like to acknowledge MIT Diagnostic Laboratory at the MIT Division of Comparative Medicine, especially E. Jordan for consultation and processing of serum samples. We would like to acknowledge MIT Department of Chemistry Instrumentation Facility for use of MALDI-ToF MS instrument.

## Table of Content graphic

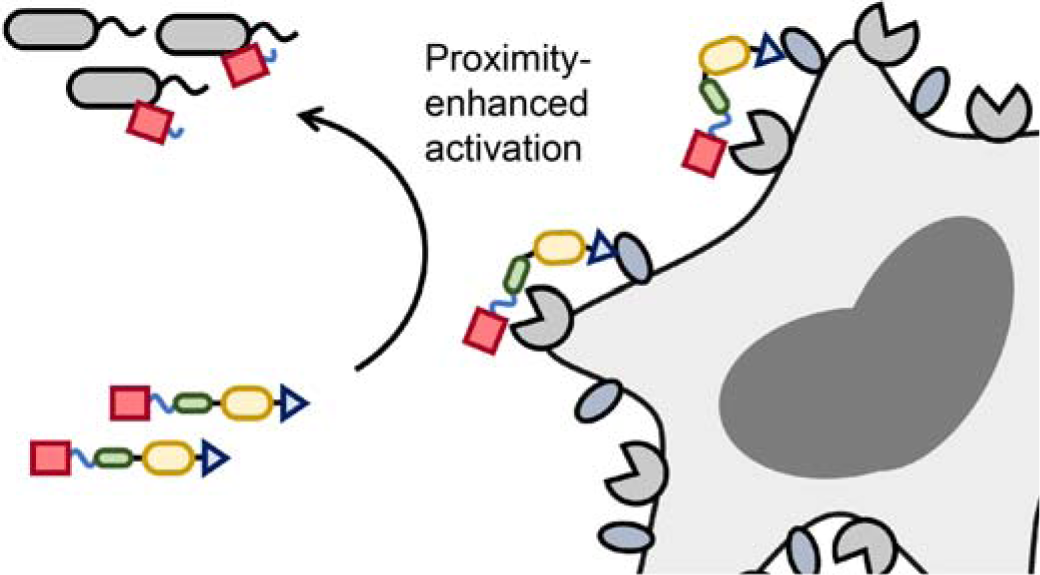

