## Supplementary information for "Nanobody-targeted conditional antimicrobial therapeutics"

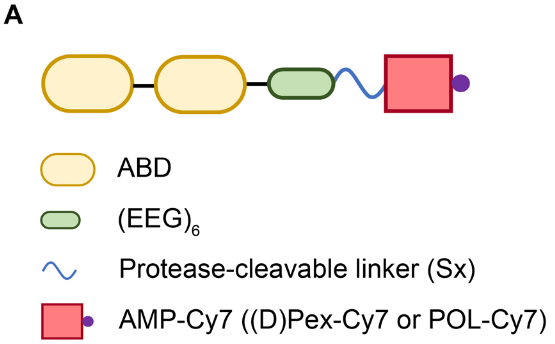


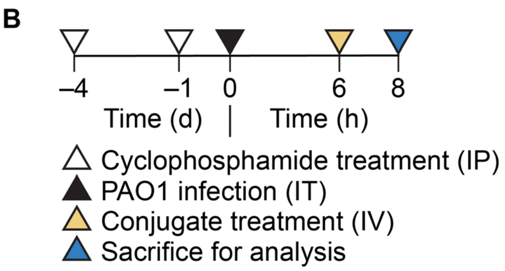


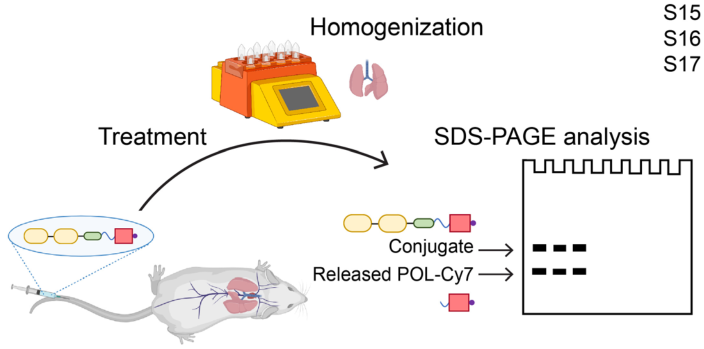


C

| Round | Substrate | Sequence | Proteases known to cleave |
|---|---|---|---|
| 1 | S1 | PLGLRSW | Thrombin, Cathepsins, MMPs |
| | S2 | KPILFFRL | Cathepsin D/E |
| | S3 | KAFRRSG | Cathepsins, Klk1 |
| | S4 | TTFYRRGA | Klk1 |
| | S5 | ARLYSRG | Klk1 |
| | S6 | TSVLMAAPQ | Napsin |
| | S7 | VGPSQG | FAP |
| 2 | S5 | ARLYSR | Klk1 |
| | S8 | VRFRST | Klk13 |
| | S9 | IQQRSL | Klks |
| | S10 | RQSRIV | Klks |
| | S11 | LAQAFRS | Klks, MMPs, ADAM10/17 |
| | S12 | TRFYSR | Klk1 |
| 3 | S11 | LAQAFRS | Klks, MMPs, ADAM10/17 |
| | S13 | LAQAVRS | ADAM10/17 |
| | S14 | LAQAFTS | ADAM17 |
| | S15 | LAAAVVS | ADAM17 |
| | S16 | KIEAVKS | ADAM10/17 |
| | S17 | PRAEALK | ADAM10/17 |

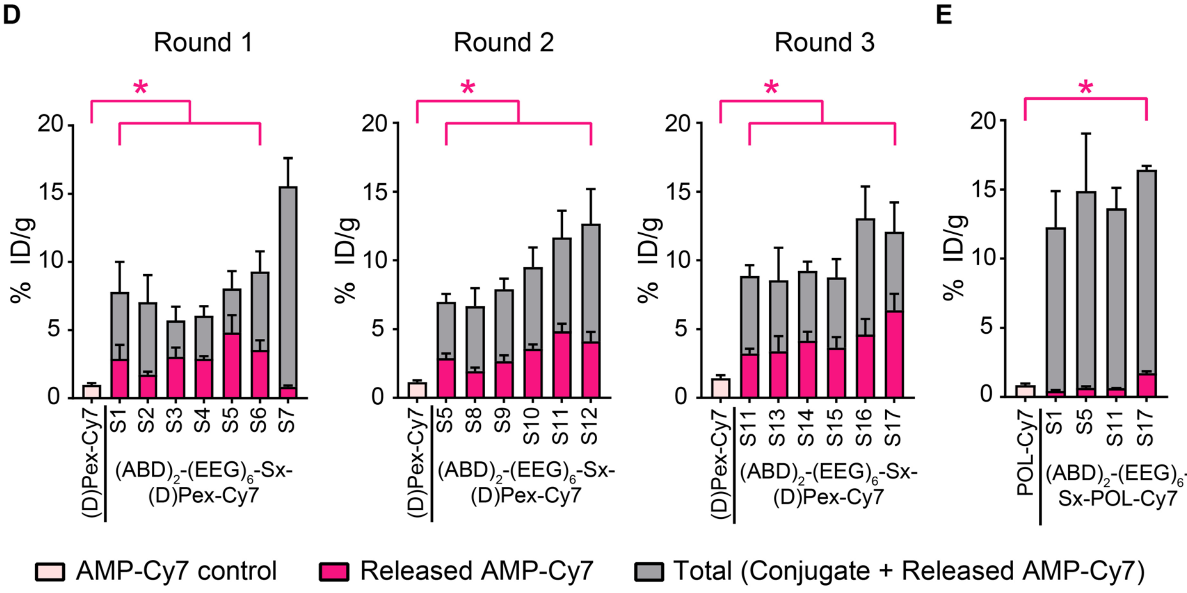

**Figure S1. Optimization of cleavable linker improves conditional therapeutic activation.**
(A) Design of ABD-AMP conjugates $(ABD)_2$-$(EEG)_6$-Sx-AMP-Cy7. (B) Experimental timeline and workflow for *in vivo* evaluation of biodistribution and activation of AMP-Cy7 conjugates. (C) List of cleavable linker substrates used in each round of screening. Quantification of total and activated fractions of (D) $(ABD)_2$-$(EEG)_6$-Sx-(D)Pex-Cy7 and (E) $(ABD)_2$-$(EEG)_6$-Sx-POL-Cy7 in PAO1-infected lungs presented as % ID/g. Panels D and E were plotted as mean ± SD and analyzed with One-way ANOVA with Tukey post hoc tests. (n = 3). Selected comparisons between AMP-Cy7 and released AMP-Cy7 from the conjugates were shown in pink. * denotes statistical significance ($P < 0.05$). Panel B was partly created with BioRender.com.

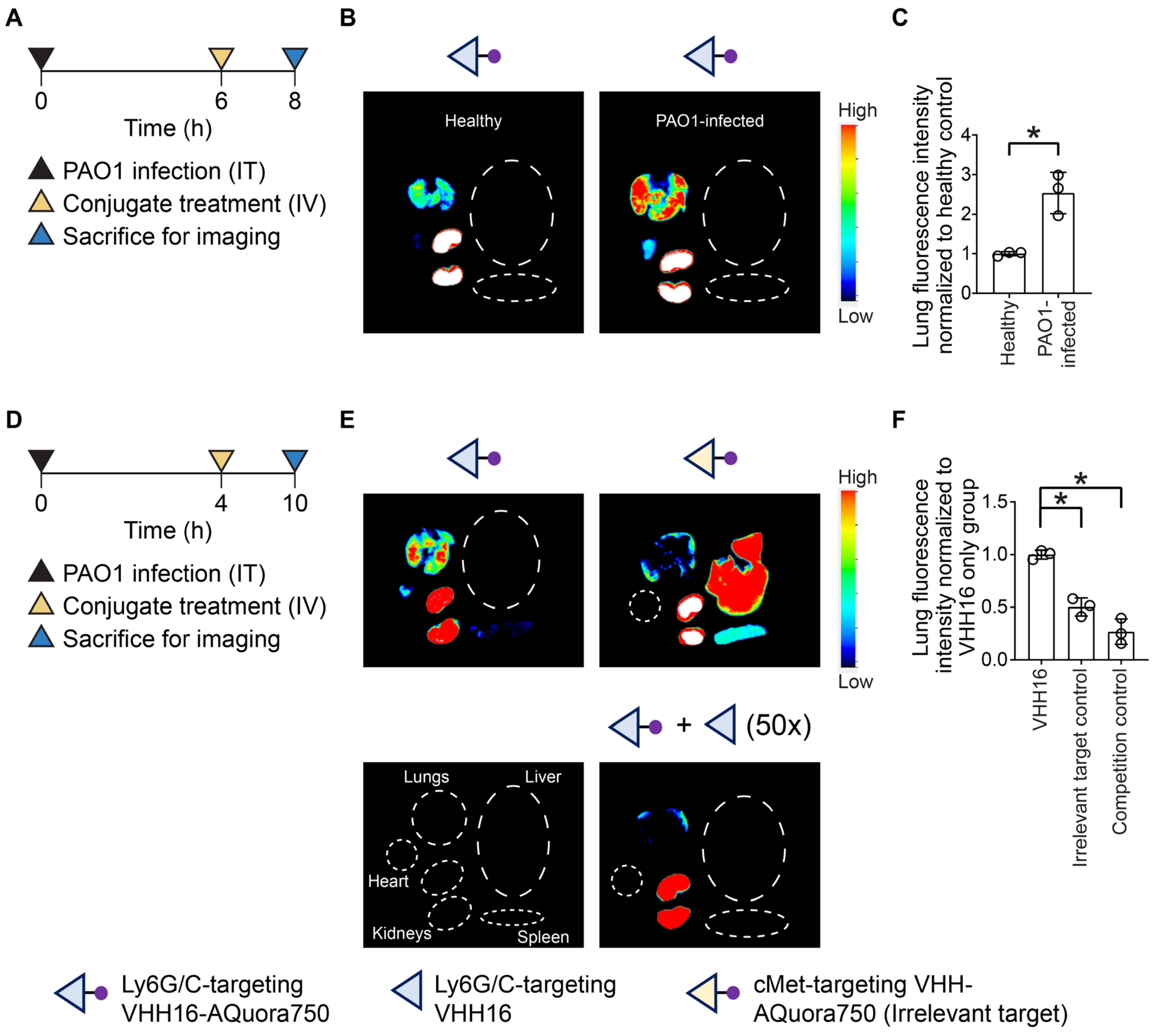


**Figure S2. VHH16 accumulated in PAO1-infected lungs in an infection-dependent, VHH-specific manner.** (A) Experimental timeline for *in vivo* biodistribution evaluation of VHH16-AQuora750 in healthy and PAO1-infected mice. (B) Representative *ex vivo* fluorescence images of VHH16-AQuora750 accumulation in different organs. (C) Quantification of VHH16-AQuora750 accumulation in healthy and PAO1-infected lungs reported as fluorescence intensity normalized to that of the healthy control. (D) Experimental timeline for *in vivo* biodistribution evaluation of VHH16-AQuora750, cMet-targeting VHH-AQuora750 (Irrelevant target control) and VHH16-AQuora750 + excess VHH16 (Competition control) in PAO1-infected mice. (E) Representative *ex vivo* fluorescence images of VHH-AQuora750 accumulation in different organs. (F) Quantification of VHH-AQuora750 accumulation in PAO1-infected lungs reported as fluorescence intensity normalized to that of the VHH16 only control. Panels C and F were plotted as mean ± SD and analyzed with One-way ANOVA with Tukey post hoc tests. (n = 3). * denotes statistical significance ($P < 0.05$).

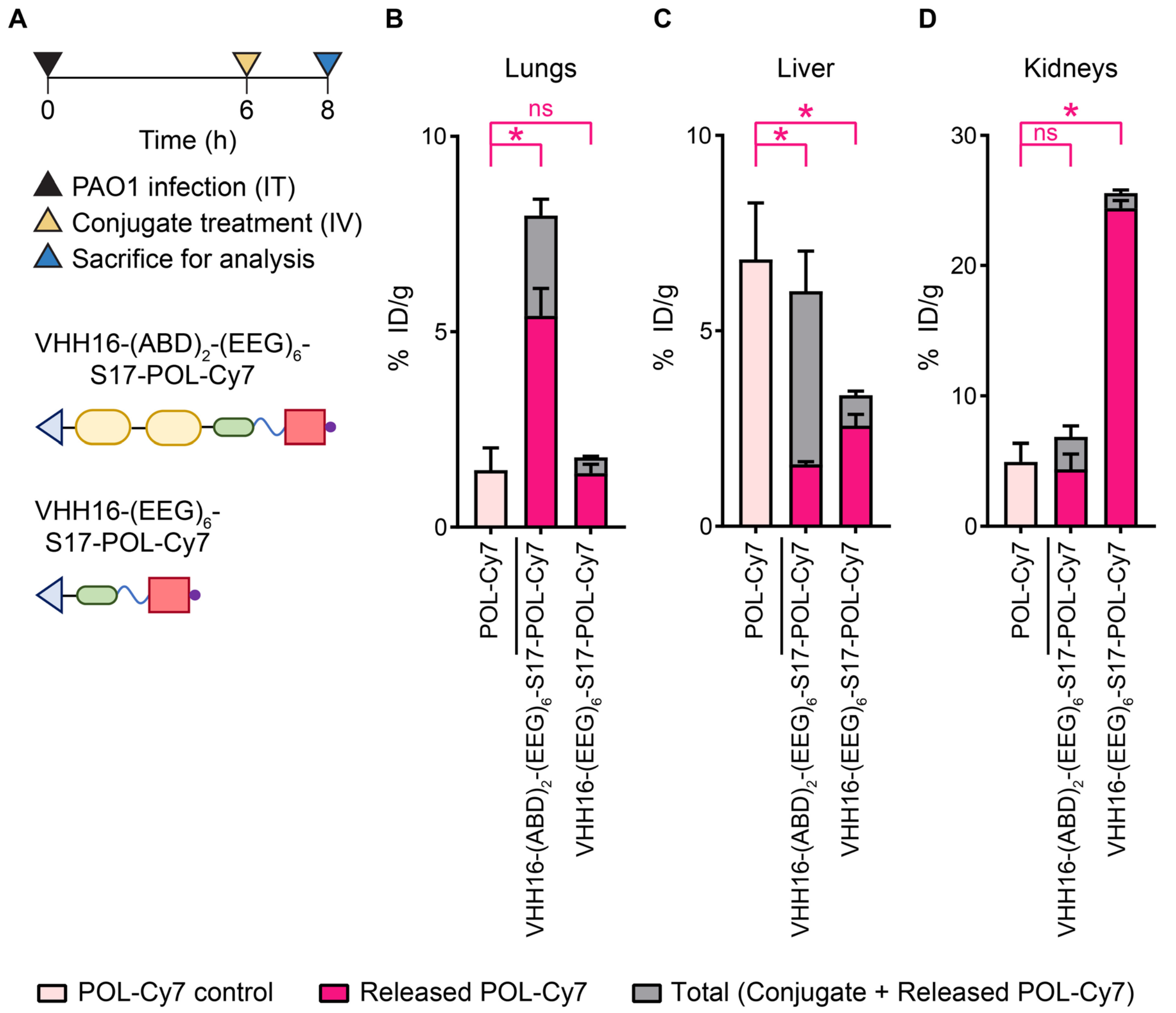


**Figure S3. ABD is required to increase the amount of released POL-Cy7 of VHH16-targeted conjugate in PAO1-infected lungs.**

(A) Experimental timeline for *in vivo* evaluation of biodistribution and activation of VHH16-targeted POL-Cy7 conjugates with and without ABD. Quantification of total and activated fractions of the POL-Cy7 conjugates in (B) PAO1-infected lungs, (C) liver, and (D) kidneys presented as % ID/g. Panels B-D were plotted as mean ± SD and analyzed with One-way ANOVA with Tukey post hoc tests. (n = 3). Comparisons between POL-Cy7 and released POL-Cy7 from the conjugates were shown in pink. * denotes statistical significance ($P < 0.05$).

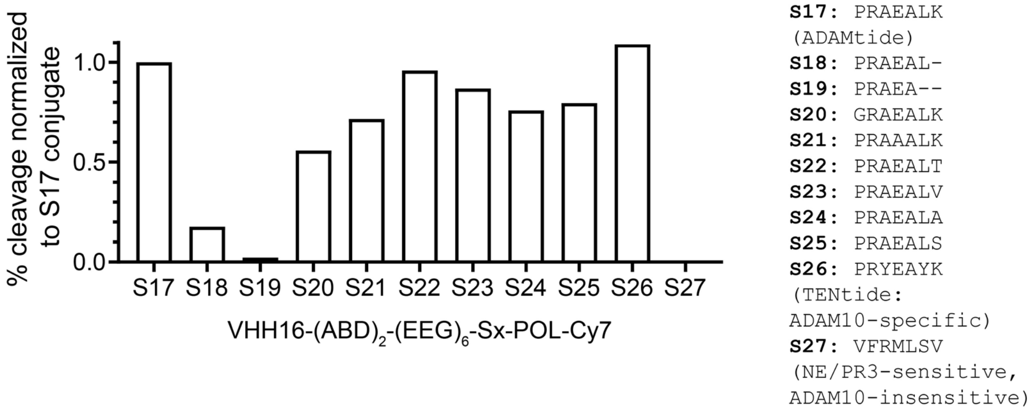


**Figure S4.** ***In vitro*** **ADAM10 cleavage assay identified tolerable mutations of the S17 linker.**
*In vitro* cleavages of VHH16-(ABD)$_2$-(EEG)$_6$-Sx-POL-Cy7 with different cleavable linkers by human ADAM10 were determined by SDS-PAGE analysis after incubation for 24 h and plotted as % cleavage normalized to that of the S17 conjugate.

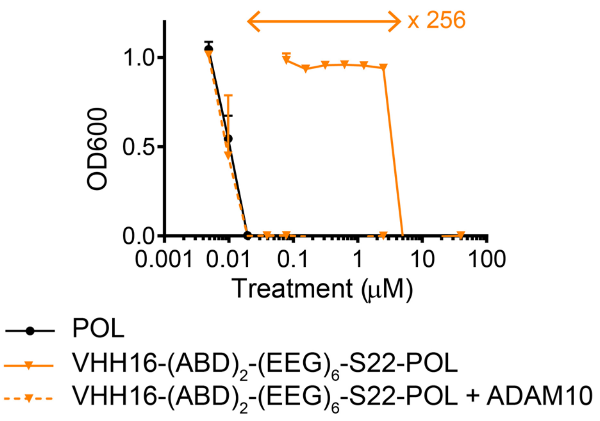


**Figure S5. Activated VHH16-(ABD)$_2$-(EEG)$_6$-S22-POL has an equivalent antimicrobial potency as free POL.** *In vitro* evaluation of antimicrobial activity masking of VHH16-(ABD)$_2$-(EEG)$_6$-S22-POL via microdilution assay on PAO1. Bacterial viabilities were measured based on OD600 absorbance normalized to the untreated control and plotted as mean ± SD. (n = 3).

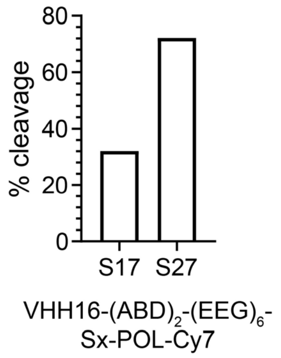


**Figure S6. S17 and S27 POL-Cy7 conjugates can be activated to neutrophil elastase.**
*In vitro* cleavages of VHH16-$(ABD)_2$-$(EEG)_6$-Sx-POL-Cy7 with different cleavable linkers by human neutrophil elastase (NE) were determined by SDS-PAGE analysis after incubation for 2 h and plotted as % cleavage.

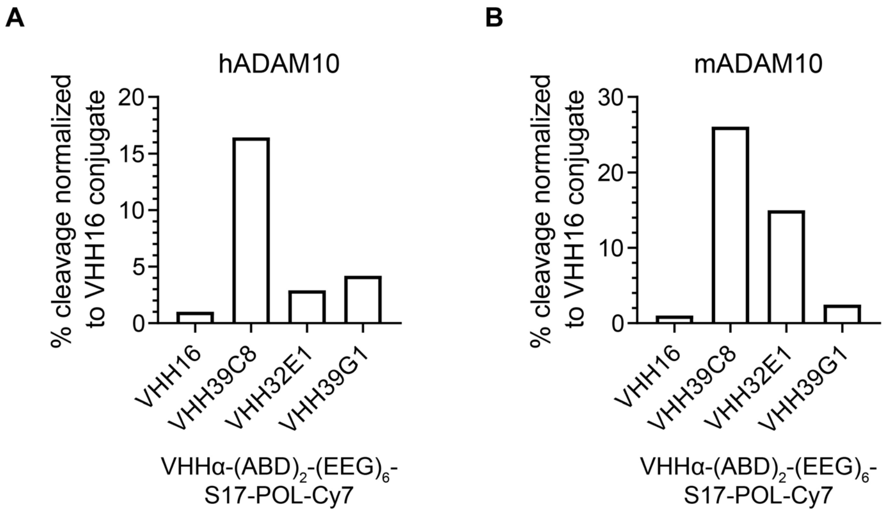


**Figure S7. ADAM10-targeting VHHs enhance *in vitro* conjugate activation by ADAM10.**
*In vitro* cleavages of VHHα-$(ABD)_2$-$(EEG)_6$-S17-POL-Cy7 with different ADAM10-targeting VHHs by human ADAM10 (left) and mouse ADAM10 (right) were determined by SDS-PAGE analysis after incubation for 2 h and plotted as % cleavage normalized to that of the Ly6G/C-targeted VHH16 control conjugate.

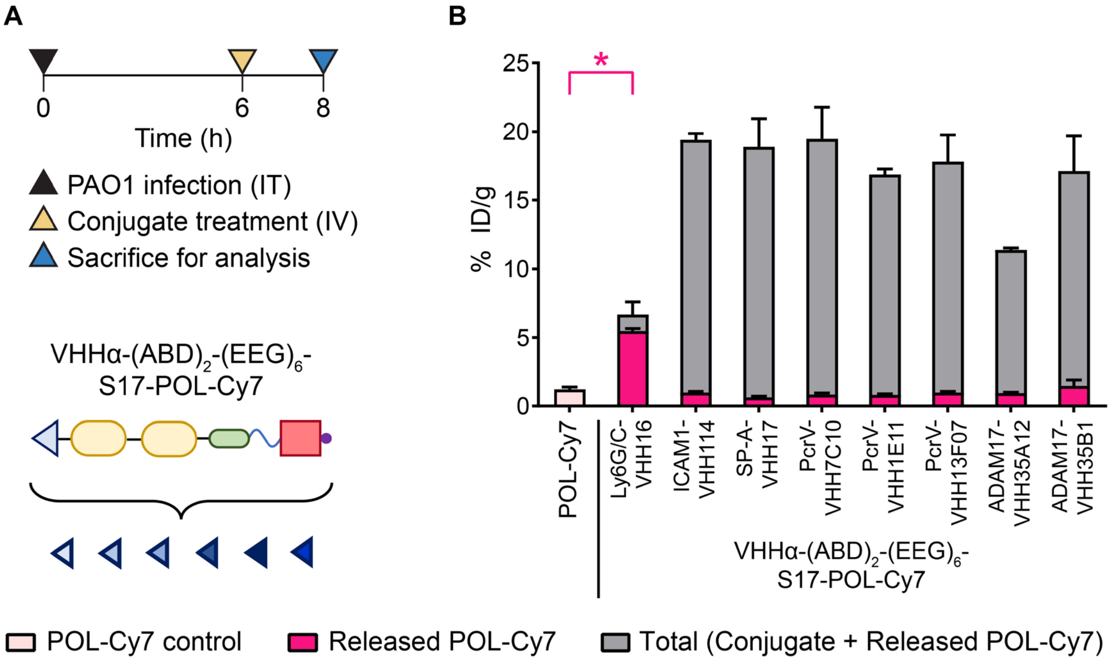


**Figure S8. Expanded screening of VHHs with relevant targets for enhanced conjugate activation.** (A) Experimental timeline for *in vivo* evaluation of biodistribution and activation of VHHα-$(ABD)_2$-$(EEG)_6$-S17-POL-Cy7 with different targeting VHHs (VHHα). (B) Quantification of total and activated fractions of the POL-Cy7 conjugates in PAO1-infected lungs. Panel B was plotted as mean ± SD and analyzed with One-way ANOVA with Tukey post hoc tests. (n = 3). Only the amount of released POL-Cy7 from the VHH16 conjugate is statistically higher than that of the POL-Cy7 control indicated in pink asterisk. * denotes statistical significance ($P < 0.05$).

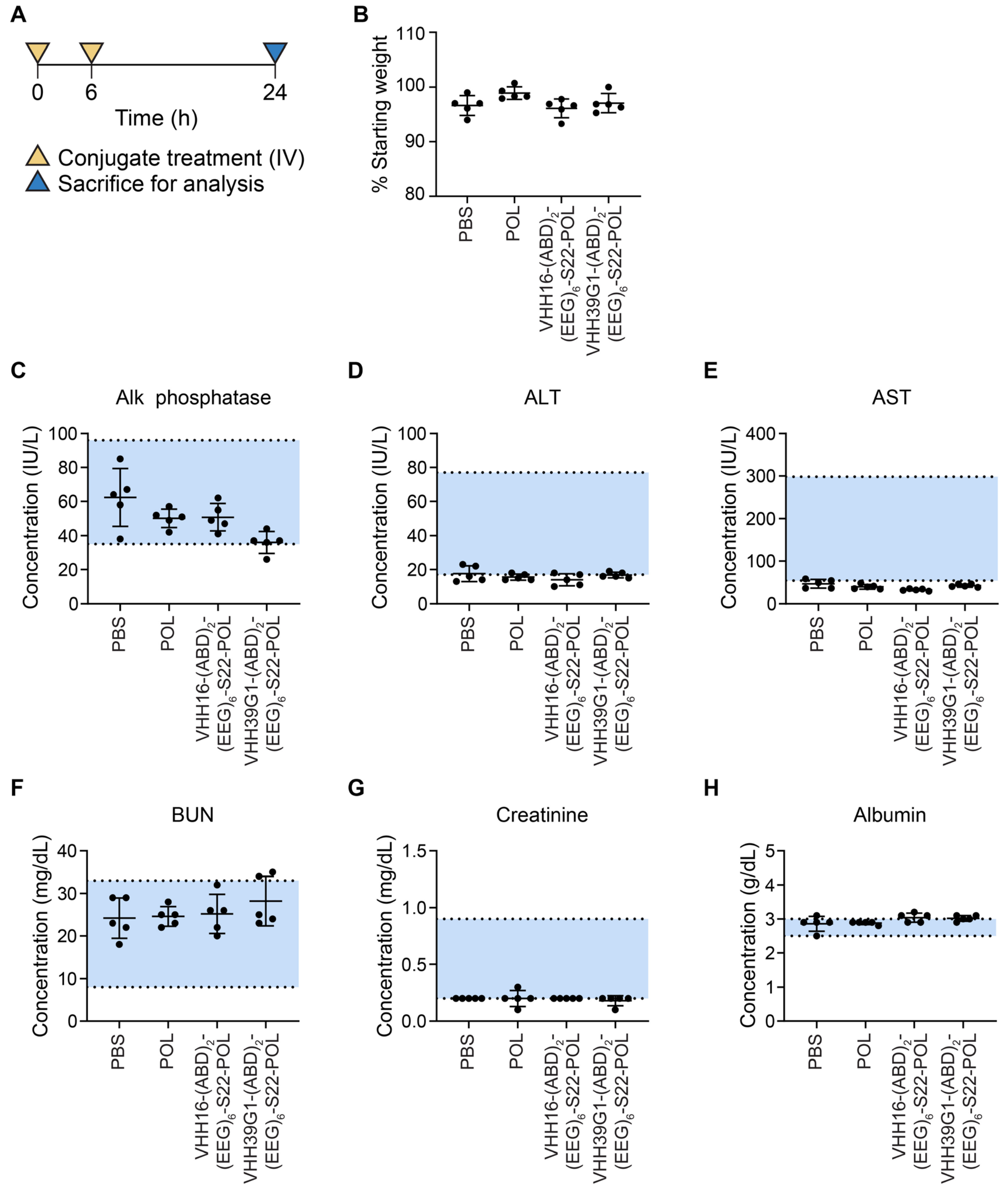


**Figure S9. VHH-targeted POL conjugates exhibit good safety profiles.**

(A) Experimental timeline for *in vivo* toxicity evaluation of VHH16-$(ABD)_2$-$(EEG)_6$-S22-POL and VHH39G1-$(ABD)_2$-$(EEG)_6$-S22-POL. Mice were treated intravenously with the conjugates at 5 mg/kg POL eq. twice (Total of 10 mg/kg POL eq. dose) and sacrificed at 24 h post first treatment to collect serums. (B) Percent body weight at the end point relative to the starting weight. Serum analysis of (C) Alkaline (Alk) phosphatase, (D) Alanine aminotransferase (ALT), (E) Aspartate aminotransferase (AST), (F) Blood urea nitrogen (BUN), (G) Creatinine, and (H) Albumin. Panels B-H was plotted as mean ± SD. (n = 5). Blue area indicates a normal reference range.

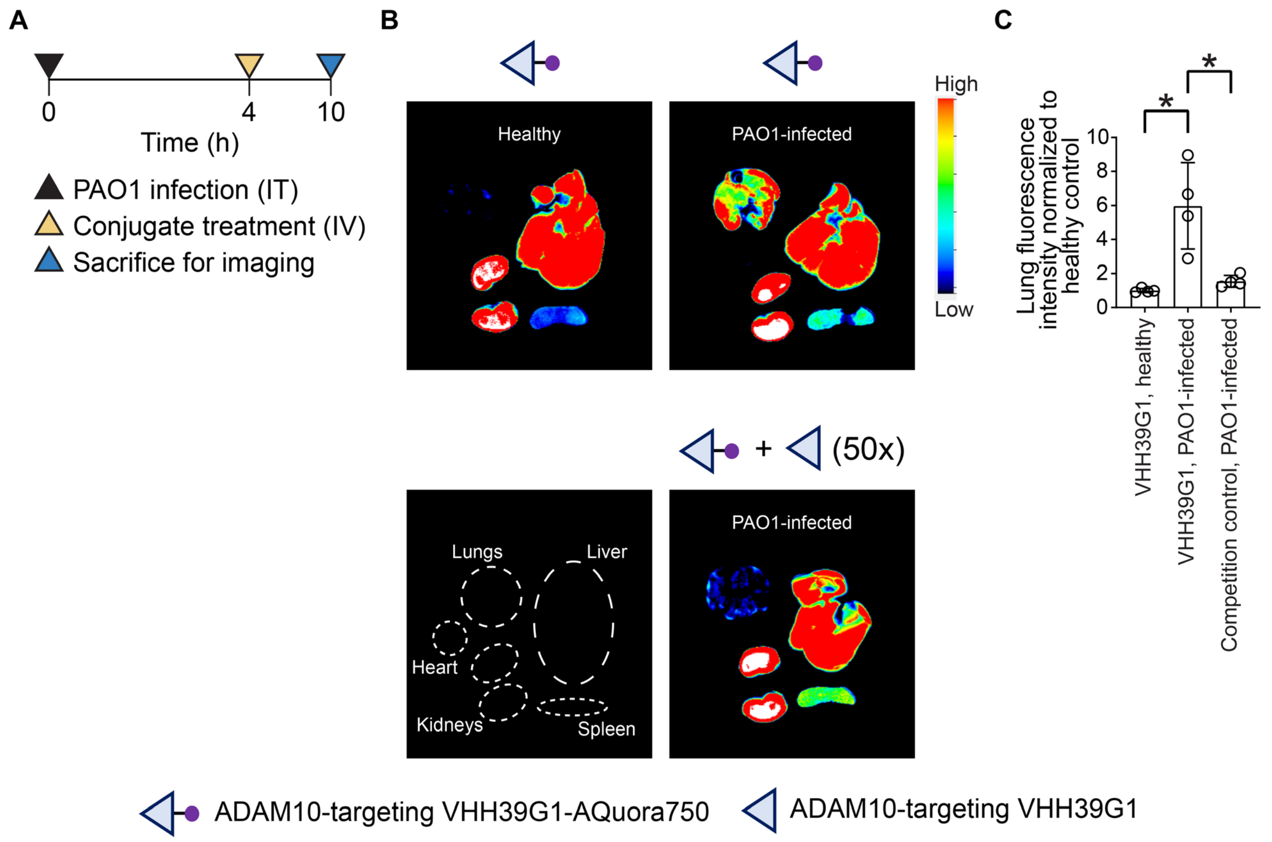


**Figure S10. VHH39G1 accumulated in PAO1-infected lungs in an infection-dependent, VHH-specific manner.**

(A) Experimental timeline for *in vivo* biodistribution evaluation of VHH39G1-AQuora750 in healthy and PAO1-infected mice without and with excess VHH39G1 (Competition control). (B) Representative *ex vivo* fluorescence images of VHH39G1-AQuora750 accumulation in different organs. (C) Quantification of VHH39G1-AQuora750 accumulation in healthy and PAO1-infected lungs without and with competition with excess VHH39G1 reported as fluorescence intensity normalized to that of the healthy control. Panel C was plotted as mean ± SD and analyzed with One-way ANOVA with Tukey post hoc tests. (n = 4). * denotes statistical significance ($P < 0.05$).

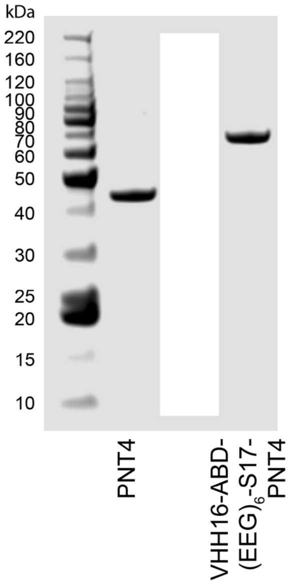


**Figure S11. Recombinant VHH16-ABD-(EEG)$_6$-S17-PNT4 was readily expressed.**
SDS-PAGE analysis of recombinantly expressed PNT4 and VHH16-ABD-(EEG)$_6$-S17-PNT4.

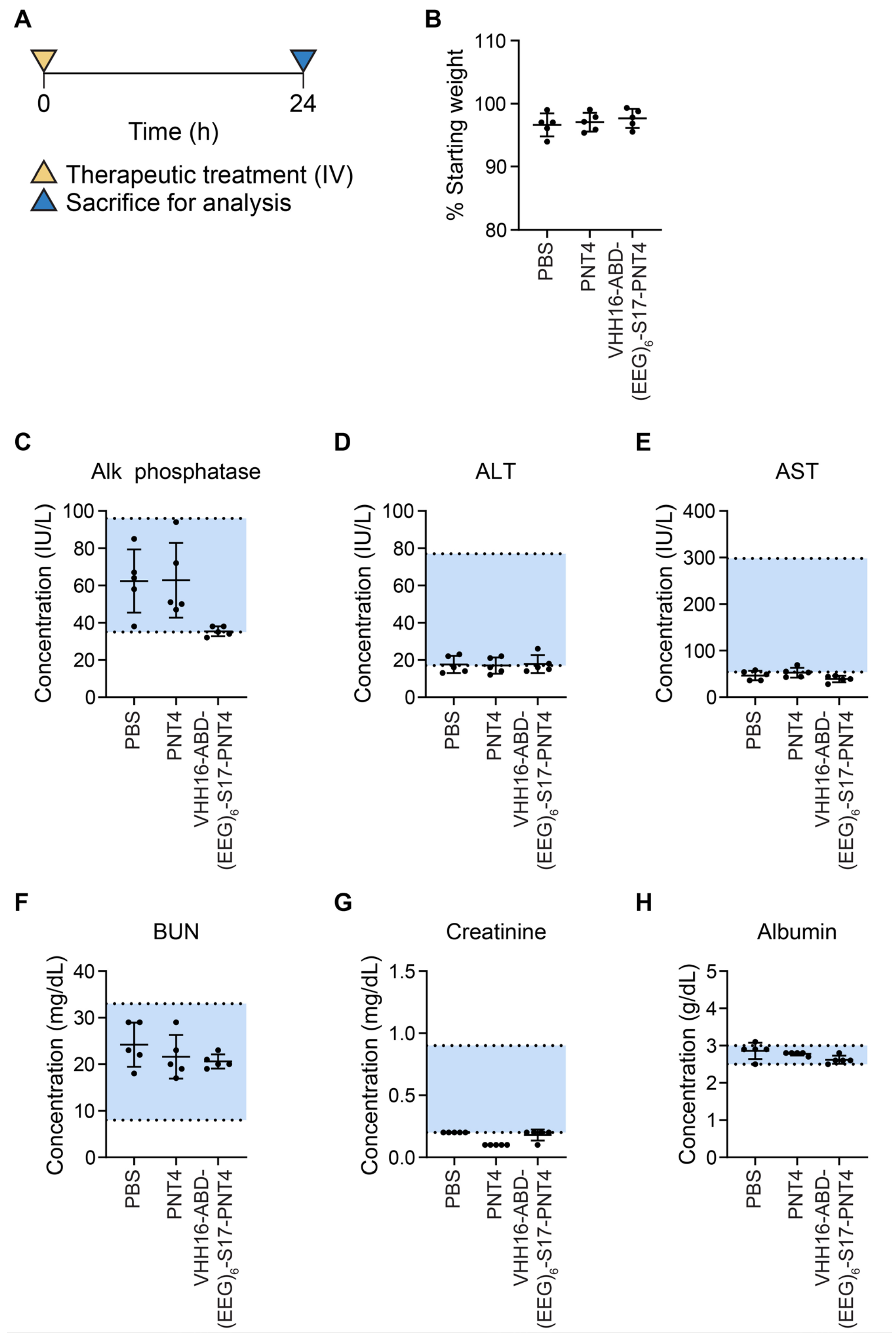


**Figure S12. VHH-targeted conditional PNT4 exhibits a good safety profile.**

(A) Experimental timeline for *in vivo* toxicity evaluation of VHH16-ABD-$(EEG)_6$-S17-PNT4. Mice were treated intravenously with VHH16-ABD-$(EEG)_6$-S17-PNT4 at 5 mg/kg PNT4 eq. and sacrificed at 24 h post treatment to collect serums. (B) Percent body weight at the end point relative to the starting weight. Serum analysis of (C) Alk phosphatase, (D) ALT, (E) AST, (F) BUN, (G) Creatinine, and (H) Albumin. Panels B-H was plotted as mean ± SD. (n = 5). Blue area indicates a normal reference range.

**Table S1. List of therapeutic peptides and sequences.**

| Peptide | Sequence (N→C) |
|---|---|
| (D)Pex-Cy7-azide | Azidoacetyl–GiGkflkkakkfGkafvkilkk–K(Cy7)–CONH2 |
| POL | Cyclo-(T-W-I-(Dab)-(Orn)-(Dab)-(Dab)-W-(Dab)-(Dab)-A-S-p-P) |
| POL-Cy7 | Cyclo-(K(Cy7)-W-I-(Dab)-(Orn)-(Dab)-(Dab)-W-(Dab)-(Dab)-A-S-p-P) |
| POL-Cy7-azide | Cyclo-(K(N3)-W-I-(Dab)-(Orn)-(Dab)-(Dab)-W-(Dab)-(Dab)-K(Cy7)-S-p-P) |
| Azido-S17-POL | Cyclo-(K(Azidoacetyl-PEG2-PEG2-GPRAEALK-)-W-I-(Dab)-(Orn)-(Dab)-(Dab)-W-(Dab)-(Dab)-A-S-p-P)<br>Azidoacetyl-PEG2-PEG2-GPRAEALK was grafted from the Lys side chain (Highlighted in red) |
| Azido-S22-POL | Cyclo-(K(Azidoacetyl-PEG2-PEG2-GPRAEALT-)-W-I-(Dab)-(Orn)-(Dab)-(Dab)-W-(Dab)-(Dab)-A-S-p-P)<br>Azidoacetyl-PEG2-PEG2-GPRAEALT was grafted from the Lys side chain (Highlighted in red) |
| Azido-S28-POL | Cyclo-(T-W-I-(Dab)-(Orn(Azidoacetyl-G-Nle(O-Bzl)–Met(O)2–Oic–Abu-))-(Dab)-(Dab)-W-(Dab)-(Dab)-A-S-p-P)<br>Azidoacetyl-G-Nle(O-Bzl)–Met(O)2–Oic–Abu was grafted from the Orn side chain (Highlighted in red) |

Note:
Cyclo indicates head-to-tail lactam cyclization.
Small letters denote D-amino acid.

Dab = 2,4-diaminobutyric acid
Orn = L-ornithine
Abu = L-2-aminobutyric acid
Nle(O-Bzl) = 6-benzyloxy-L-norleucine
Met(O)2 = L-methionine sulfone
Oic = Octahydroindole-2-carboxylic Acid
PEG2 = 2-(2-(2-Aminoethoxy)ethoxy)acetic acid

For the non-labeled therapeutic peptide conjugates, the cleavable linkers were synthesized in fusion to the therapeutic peptide to minimize the cleavage scar.

For the Cy7-labeled therapeutic peptide conjugates, the cleavable linkers were encoded in the recombinantly expressed fusion protein carrier VHHα-$(ABD)_2$-$(EEG)_6$-Sx-GC before DBCO-Mal functionalization and conjugation with AMP-Cy7-azide.

**Table S2. List of VHH clones.**

| Target | Clone |
|---|---|
| Ly6G/C | VHH16, VHH21[1] |
| CD11b | VHH13[2] |
| ADAM10 | VHH39C8, VHH32E1, VHH39G1[3] |
| ADAM17 | VHH35A12, VHH35B1[3] |
| ICAM-1 | VHH11-4[4] |
| SP-A | VHH17[5] |
| PcrV | VHH7C10, VHH1E11, 13F07[6] |
